# Removal of *pomt1* in zebrafish leads to loss of α-dystroglycan glycosylation and dystroglycanopathy phenotypes

**DOI:** 10.1101/2022.10.15.512359

**Authors:** Brittany F. Karas, Kristin R. Terez, Shorbon Mowla, Namarata Battula, Kyle P. Flannery, Brian M. Gural, Grace Aboussleman, Numa Mubin, M. Chiara Manzini

## Abstract

Biallelic mutations in *Protein O-mannosyltransferase 1* (*POMT1*) are among the most common causes of a severe group of congenital muscular dystrophies (CMDs) known as dystroglycanopathies. POMT1 is a glycosyltransferase responsible for the attachment of a functional glycan mediating interactions between the transmembrane glycoprotein dystroglycan and its binding partners in the extracellular matrix (ECM). Disruptions in these cell-ECM interactions lead to multiple developmental defects causing brain and eye malformations in addition to CMD. Removing *Pomt1* in the mouse leads to early embryonic death due to the essential role of dystroglycan during placental formation in rodents. Here, we characterized and validated a model of *pomt1* loss of function in the zebrafish showing that developmental defects found in individuals affected by dystroglycanopathies can be recapitulated in the fish. We also discovered that *pomt1* mRNA provided by the mother in the oocyte supports dystroglycan glycosylation during the first few weeks of development. Muscle disease, retinal synapse formation deficits, and axon guidance defects can only be uncovered during the first week post fertilization by generating knock-out embryos from knock-out mothers. Conversely, maternal *pomt1* from heterozygous mothers was sufficient to sustain muscle, eye, and brain development only leading to loss of photoreceptor synapses at 30 days post fertilization. Our findings show that it is important to define the contribution of maternal mRNA while developing zebrafish models of dystroglycanopathies and that offspring generated from heterozygous and knock-out mothers can be used to differentiate the role of dystroglycan glycosylation in tissue formation and maintenance.

## Introduction

Dystroglycanopathies are a group of rare autosomal recessive neuromuscular disorders which include the most severe forms of congenital muscular dystrophy (CMD). These diseases also affect the eye and brain and lead to early mortality (1,2). The primary molecular deficit is the loss of interactions between the extracellular portion of the transmembrane glycoprotein dystroglycan, a -dystroglycan (a -DG), and ligands in the extracellular matrix (ECM) (2). These interactions are mediated via an elongated, specialized O-linked glycan on a-DG termed matriglycan assembled via multiple glycosyltransferases (2,3). Biallelic mutations in *dystroglycan* (*DAG1*, OMIM:128239) itself have been identified as the cause of primary dystroglycanopathy (OMIM:613818, 616538), but they are exceedingly rare (4). Most cases of dystroglycanopathies are deemed secondary and are caused by variants in the glycosyltransferases involved in catalyzing the synthesis of glycan chain in the endoplasmic reticulum (ER) and Golgi (2). Lastly, tertiary dystroglycanopathies are caused by mutations in genes that either control glycosyl donor production or a-DG trafficking (5).

*Protein O-mannosyltransferase 1* (*POMT1*, OMIM:607423) catalyzes the addition of an O-linked mannose to a -DG starting the assembly of the functional glycan as the protein is translated in the ER (6). *POMT1* mutations are one of the most frequent causes of dystroglycanopathy in multiple populations and lead to the full spectrum of disease from limb-girdle muscular dystrophy (POMT1-LGMD, OMIM:609308) to severe CMD disorders affecting the brain and the eyes such as Walker Warburg Syndrome (WWS) (OMIM:236670) (7–9). Global knock-out (KO) of *Pomt1* in the mouse leads to early embryonic lethality similarly to *Dag1* mutants. This is due to the critical role of dystroglycan in Reichert’s membrane, a specialized basement membrane in rodent embryos (10,11). Thus, *Pomt1* mouse models cannot fully recapitulate the disease condition and have only been studied using conditional approaches (12,13).

We sought to develop a novel animal model of dystroglycanopathies by characterizing *pomt1* loss of function in the zebrafish. The zebrafish has become an established model for muscle disease because of the molecular and physiological conservation of disease mechanisms, their external embryonic development, and their fecundity accompanied with small size which allows for high-throughput drug screening for motor phenotypes (14,15). Most zebrafish models for dystroglycanopathies to date have been generated via gene knockdown using morpholino oligonucleotides (MOs) to validate genetic findings in humans. The same array of phenotypes including loss of a-DG glycosylation, shortened body axis, reduced mobility, muscular dystrophy, and eye and brain malformation were found in knockdown models of multiple glycosyltransferases (16–20). However, knockdown approaches are limited by their variable and transient effect. Furthermore, results of *pomt1* knockdown in larvae were unexpectedly mild compared to other dystroglycanopathy zebrafish knockdowns (19).

Out of the nineteen dystroglycanopathy genes, there are currently four zebrafish genetic models reported. The *patchytail* mutant was generated via a mutagenesis screen leading to a missense mutation in *dag1* and loss of the protein. These mutants die by 10 days post fertilization and show similar deficits to the morphants, although less severe, which recapitulate multiple aspects of the human presentation in the muscle, eye, and brain (21). KO larvae for the glycosyltransferase *fukutin-related protein* (*fkrp*) more closely phenocopied the morphants and were then used to study the effects of overexpression of the most common *FKRP* (OMIM: 606596) missense variant leading to *FKRP*-related LGMD (OMIM:607155) to identify compounds that could ameliorate motor phenotypes by drug screening (22). Interestingly, no motor phenotypes were described for zebrafish CRISPR mutants developed for glycosyltransferase *protein O-mannoseβ-1,2-N-acetylglucosaminyltransferase* (*pomgnt1*) or *Protein O-mannosyltransferase 2* (*pomt2*) (23,24). Both KOs showed photoreceptor degeneration in the retina at 6 months of age, reflecting retinal deficits identified in patients (25). Hypomorphic variants in *POMGNT1* have also been linked with non-syndromic retinitis pigmentosa (26). In addition, the *pomt2* mutant showed mild hydrocephalus and muscular dystrophy at 2 months (24). While these studies indicated that relevant phenotypes resulting from global KOs of dystroglycanopathy genes can be successfully modeled in zebrafish, these were less severe than expected in some cases. However, it remains unclear whether these differences are due to genetic compensation.

Here, we show that *pomt1* KO zebrafish show varying degree of muscle, eye, and brain phenotypes depending on whether *pomt1* mRNA is provided to the oocyte from the mother. Since the oocytes are externally fertilized, zebrafish females provide nutrients, mRNAs, and proteins to support early development in the yolk (27). *pomt1* KOs obtained from heterozygous mothers can compensate for the early developmental phenotypes by having maternally provided Pomt1 glycosylate a-DG. More severe phenotypes appear when KO embryos are generated by a KO mother. Our work indicates that special consideration must be taken regarding the potential maternal contribution in future dystroglycanopathy mutants.

## Results

### A late nonsense allele in pomt1 leads to complete protein loss and loss of α-DG glycosylation

In order to determine whether loss of *pomt1* would lead to dystroglycanopathy phenotypes in the zebrafish, we obtained a *pomt1* line generated through a N-ethyl-N-nitrosurea (ENU) mutagenesis screen by the Zebrafish Mutation Project (ZMP line: sa14667, kind gift of Dr. James Dowling, Sick Kids, Toronto, Canada) (**Fig. 1A**) (28). This line introduced a variant in exon 19 of *pomt1* (NM_001048067: c.1911T>A) leading to a stop codon (NP_001041532: p.Tyr637Ter) towards the C-terminus of the 720 amino acid protein. Since multiple homozygous nonsense and frameshift variants had been reported in humans to cause the most severe form of dystroglycanopathy, WWS (29,30), we hypothesized that this zebrafish variant could lead to nonsense mediated decay of the mRNA and complete protein loss. No compensation was expected via genome duplication since *pomt1* is not duplicated in the zebrafish. We bred the p.Tyr637Ter variant into homozygosity and confirmed the variant by Sanger sequencing (**Fig. 1B**). We first tested for possible non-sense mediated decay of the mRNA through quantitative PCR (qPCR) analysis and found that *pomt1* mRNA levels were not significantly altered between 5 and 40 days post fertilization (dpf) (**Fig. 1C**). The Pomt1 protein was completely lost at 30 dpf in homozygous animals and no additional truncated band was noted, apart from a non-specific band which is also present in wild-type (WT) samples (**Fig. 1D**). We validated the Pomt1 antibody for immunohistochemistry by confirming colocalization with the sarcoplasmic reticulum (SR) marker, Ryanodine Receptor 1 (Ryr1) (**Suppl. Fig. 1A**), and also showed that Pomt1 was already absent in muscle fibers at 10 dpf (**Suppl. Fig. 1B**). In parallel, we confirmed that Pomt1 function was completely lost. No a-DG glycosylation was found using a glyco-specific antibody in wheat germ agglutinin (WGA) enriched zebrafish head tissue from 30 dpf animals, though α-DG expression was preserved (**Fig. 1E**). In summary, the *pomt1^Y637X^* line leads to complete loss of function of Pomt1 and will be termed as *pomt1* KO^Het^, to indicated it was obtained from a heterozygous cross.

**Figure 1.**
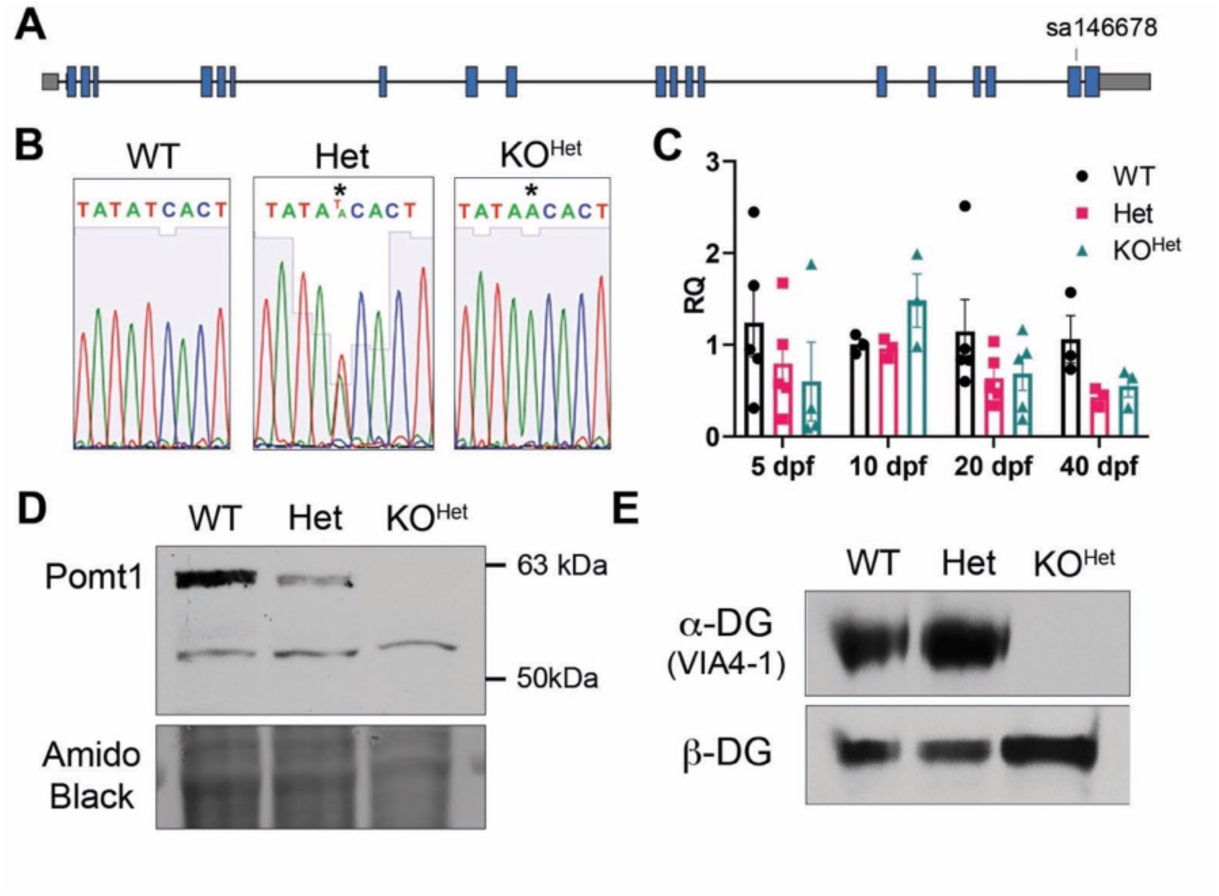
*pomt1* nonsense variant leads to complete loss of protein and protein function. **A.** Schematic of the gene structure of *pomt1* including the location of the variant in exon 19. **B.** Sanger sequencing validation shows the stop codon generated in the KO genome. **C.** qPCR analysis showed consistent reduction in *pomt1* mRNA apart from 10 dpf. All groups included 3-5 samples generated from composites of at least three animals. Although consistently trending within and over time points, data was not significant between genotypes. **D.** Pomt1 is completely absent on Western blot in 30 dpf protein lysates from KO^Het^ fish. **E.** a-DG glycosylation is absent in KO^Het^ tissue while β-DG expression is preserved.

### Loss of pomt1 from heterozygous crosses leads to reduced survival

No phenotypes had been reported in *pomt1* KO zebrafish larvae by the ZMP. Initial survival analysis revealed that *pomt1* KO^Het^ juveniles start dying between 30 dpf and 40 dpf (30 dpf: WT 24.3±8.0%, n=18; Het 49.0±9.5%, n=41; KO^Het^ 27.7±2.0%, n=17, N=3 clutches. 40 dpf: WT 28.0±4.0%, n=22, Het 60.3±7.5%, n=46, KO^Het^ 11.7±3.5%, n=11, >50 dpf: WT 35.6±4.2% n=32, Het 64.0±2.6% n=62, KO^Het^ 3.7±1.9% n= 2; N=3 clutches) with no survival observed beyond 70 dpf (**Fig. 2A**). While no morphological differences were noted at 5 and 20 dpf, significant differences in body length emerged at 30 dpf (WT 8.5±0.5 mm, n=18; Het 7.3±0.3 mm, n=41; KO^Het^ 6.2±0.2 mm, n=17; N=3 clutches, WT-Het p=0.039 *, WT-KO^Het^ p=0.0005 ***) (**Fig. 2B**). *pomt1* KO^Het^ juveniles were smaller and showed a delay in development with less prominent dorsal and anal fins (**Fig. 2C**). Interestingly, the surviving KO^Het^ fish at 40 dpf were not significantly smaller (WT 9.6±0.4 mm, n=22; Het 9.9±0.3 mm, n=46; KO^Het^ 8.5±0.4 mm, n=11) suggesting that larger fish survived.

**Figure 2.**
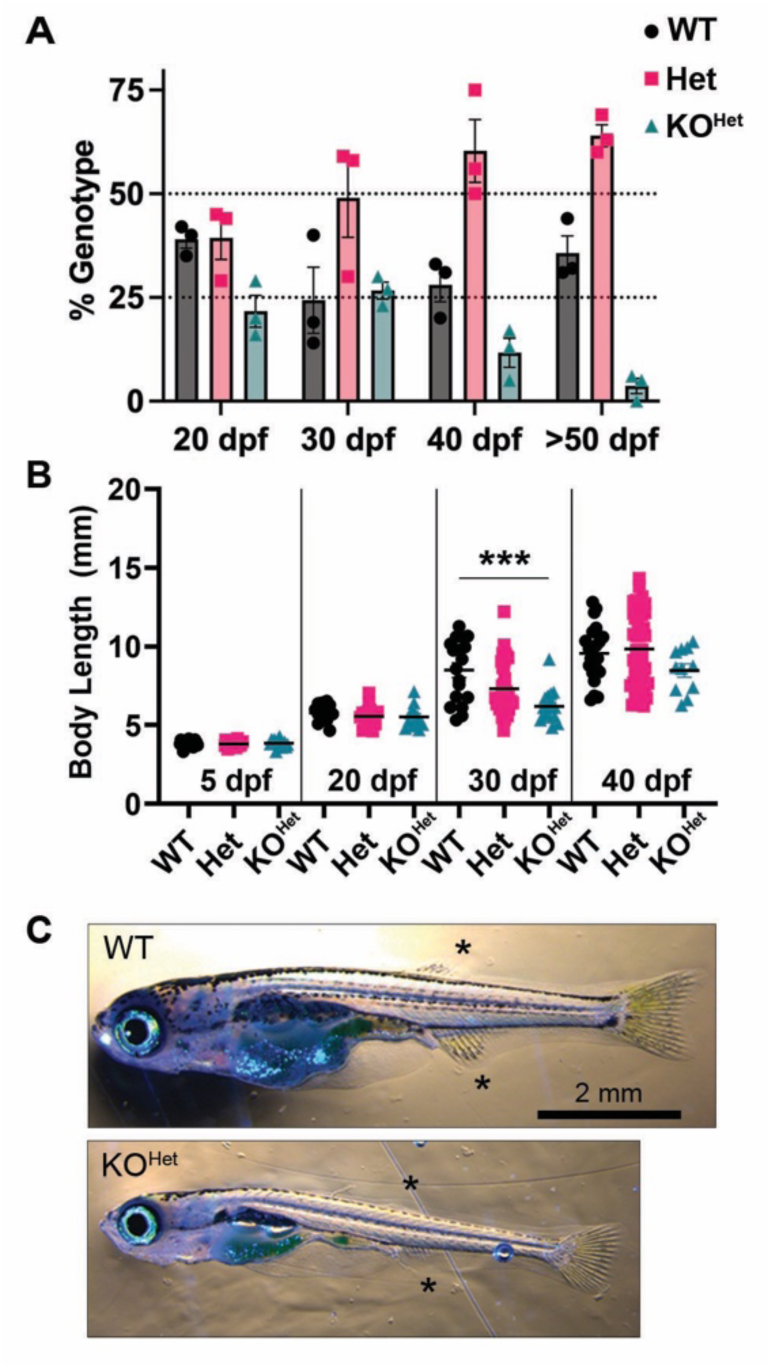
*pomt1* KO^Het^ fish show reduced survival and size. **A.** Survival is indicated as deviation from expected Mendelian ratios (25% for WTs and KO^Het^s, 50% for Hets) in 3 independent clutches showing a drop in *pomt1* KO^Het^ juveniles after 30 dpf. **B.** Significant size differences are evident in KO^Het^ fish at 30 dpf, but not in the surviving KO^Het^ fish at 40 dpf. **C.** *pomt1* KO^Het^ fish at 30 dpf appear underdeveloped with less prominent dorsal and anal fins (asterisks). Scale bar: 2mm. *** p<0.001

We tested motor function in an automated DanioVision^TM^ tracking system where the fish were allowed to swim freely for 30 minutes. No differences were noted in 5 dpf larvae (**Suppl. Fig. 2A-C**). While only a trend for reduced distance covered and swimming velocity was found in 30 dpf *pomt1* KO^Het^s (WT n=15, Het n=37, KO^Het^ n=13, N=3 clutches. Distance: WT 870.5±84.9 cm, Het 803.9±54.7 cm, KO^Het^ 666.2±88.9 cm, WT-KO^Het^ p=0.51. Velocity: WT 0.485±0.047 cm/s, Het 0.449±0.030 cm/s, KO^Het^ 0.380±0.051 cm/s, WT-KO^Het^ p=0.62) (**Fig. 3A-B**), we found a significant reduction in acceleration when compared to both Het and WT fish (WT 420.1± 24.0 cm/s^2^, Het 401.6±23.3 cm/s^2^, KO^Het^ 291.0±27.8 cm/s^2^, WT-KO^Het^ p=0.024 *, Het-KO^Het^ p=0.018 *) (**Fig. 3C**). Histological analysis of the muscle at 30 dpf was complicated by the extreme fragility of the tissue in KO^Het^ animals where myofibers tended to fragment. Via hematoxylin-eosin staining in paraffin sections, we measured myofiber length and found that this was not changed when the length was normalized for the total length of the fish (**Suppl. Fig. 2D-F**). To look at fiber size, we immunostained axial cryosections with laminin to outline the fiber boundaries and measured fiber area in a dorsal muscle at a set anatomical position (**Fig. 3D**). Though there was a shift toward larger fibers in 30 dpf KO^Het^ muscle there was not a significant change for genotype (**Fig. 3E**, also see **Suppl. Fig. 2G-H**). Although loss of *pomt1* clearly had severe effects on survival, there were no clear muscle phenotypes identified at 30 dpf which was reminiscent of the mild phenotypes identified in the *pomt2* and *pomgnt1* mutants (23,24).

**Figure 3.**
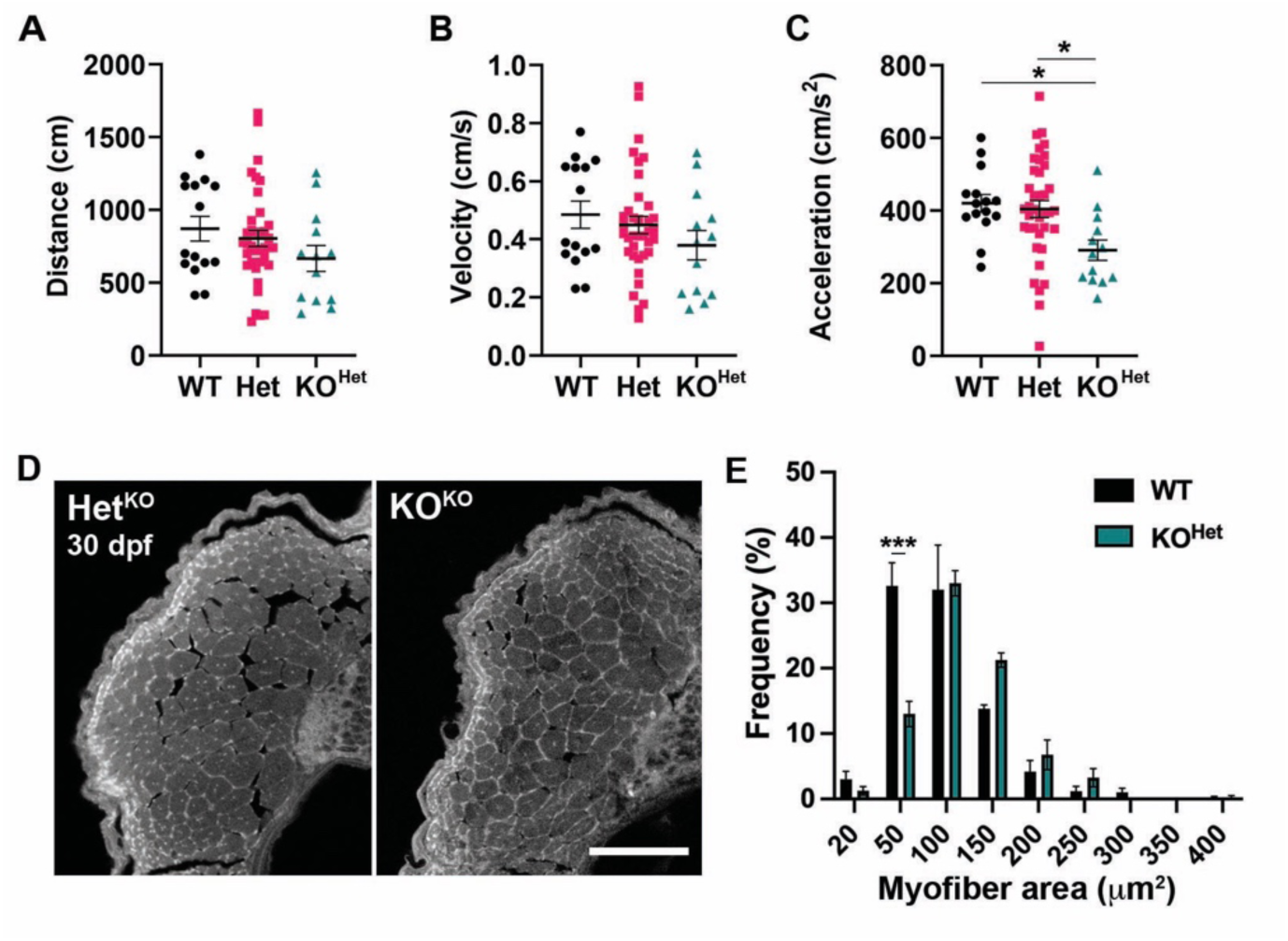
Muscle analysis in 30 dpf *pomt1* KO^Het^ juvenile zebrafish. **A-C.** Automated tracking of motor activity showed only a trend for reduction in distance (**A**) and velocity (**B**), but significant reduction in acceleration (**C**) in KO^Het^ animals compared to WT and Het. **D.** Representative dorsal muscle axial cryosections stained with laminin that were used for muscle fiber area quantification. Scale bar: 100 µm. **E.** While overall muscle fiber area was not significantly different (**Suppl. Fig. 2G**), there was shift towards larger fibers in the KO^Het^ where the frequency of fibers between 20 and 50 μm^2^ was reduced. * p<0.05

### pomt1 loss-of-function disrupts photoreceptor synapses

While the brain of 30 dpf *pomt1* KO fish showed no major structural abnormalities (**Suppl. Fig. 3**), mutant for the glycosyltransferases *pomt2* and *pomgnt1* showed photoreceptor degeneration at 6 months of age (23,24). While eye size in *pomt1* KOs was highly variable it was proportional to body size (**Suppl. Fig. 4A**). We chose to analyze the smaller *pomt1* KO^Het^ fish that were likely to be more severely affected. Retinal thickness and photoreceptor layer thickness were reduced at 30 dpf and the photoreceptor layer was disproportionally thinner than the retina (**Suppl. Fig. 4B**. Retina/photoreceptor ratio. WT n=6: 0.390±0.018, KO^Het^ n=5: 0.297±0.027, p=0.015 *). Through immunohistochemical analysis we noted several disruptions in the photoreceptor layer and in the outer plexiform layer (OPL) where photoreceptor pedicles form synapses with bipolar cells and horizontal cells begin the transmission of light signals. Immunostaining with the red/green double cone maker arrestin3a (Arr3a) which also accumulates at the presynaptic pedicles of the photoreceptors (31) showed a disruption in the organization of terminals in the OPL with some terminals retracting among the photoreceptor nuclei (**Fig. 4A**). Expression of Gnat2, a cone-specific α-transducin subunit necessary for the phototransduction cascade and color vision (32), showed reduced staining in the outer limiting membrane and in the photoreceptor outer segments (**Fig. 4A**). In addition, the rod and green cone marker, zpr-3, which is predicted to bind outer segment opsins (33) also showed localized reduced staining, though rod outer segments appear present in autofluorescence (**Fig. 4B**). OPL disorganization was evident in the Hoechst nuclear staining, where localized misalignments of photoreceptor nuclei were present, as well as smaller and misshapen horizontal cell nuclei (**Fig. 4B**). These disruptions suggested a loss of photoreceptor-bipolar cell synapses. Discontinuities in the expression of presynaptic protein synaptophysin (Syp) were present wherever photoreceptor pedicles were lost (**Fig. 4C**). Overall, our results were similar to the phenotypes in both *pomt2* and *pomgnt1* KO zebrafish and following *Pomt1* conditional removal in photoreceptors in mice where loss of dystroglycan glycosylation leads to disruptions at photoreceptor ribbon synapses and subsequent photoreceptor degeneration (13,23).

**Figure 4.**
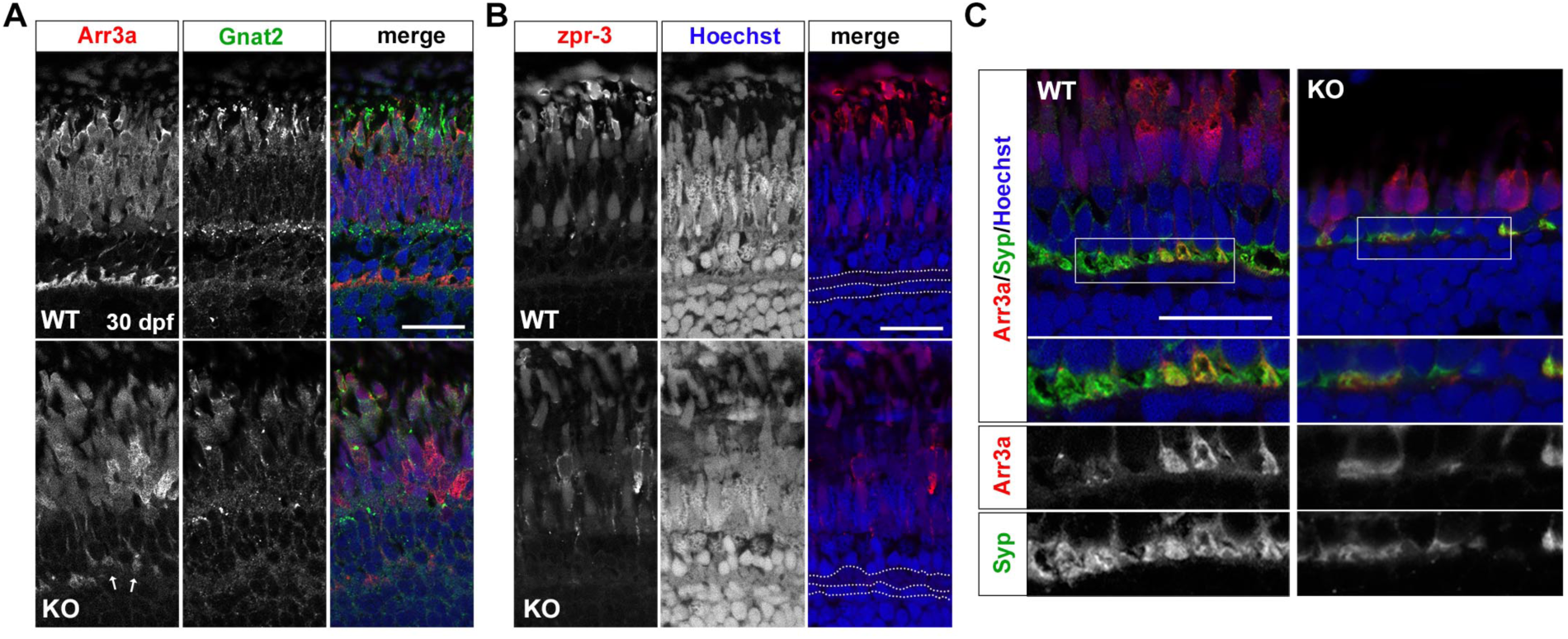
Photoreceptor synapses are disrupted in *pomt1* KO^Het^ retinae at 30 dpf. **A.** Red/green double cone marker arrestin 3a (Arr3a) shows both disorganization in the outer segment and retraction of photoreceptors pedicles in the outer plexiform layer (OPL) (arrows). Cone-specific α-transducin (Gnat2) staining is also reduced and lost in the outer limiting membrane. ONL: outer nuclear layer; INL: inner nuclear layer. Scale bar: 20 µm **B.** zpr-3 staining is reduced, but rods appear present in autofluorescence. Nuclear staining with Hoechst shows disorganization in the OPL and disruption in the nuclei of the horizontal cells below the OPL. OPL and horizontal cell nuclear layer are outlined by the white dotted lines. Scale bar: 20 µm **C.** Synaptophysin (Syp) staining is discontinuous and showing loss of synaptic contacts. The white boxes outline the inset where immunostaining for each antibody is shown below. Scale bar: 20 µm

### More severe dystroglycanopathy phenotypes are present when pomt1 is removed from fertilized oocytes

Even though eye phenotypes characteristic of dystroglycanopathies were present in the *pomt1* KO^Het^ juveniles, phenotypic presentation was later than expected when compared to *dag1* and *fkrp* zebrafish mutants (21,22). Not only are zebrafish notorious for being able to compensate for genetic mutations (34,35), but also display a fundamental difference from mammals in that a deposit of mRNA and proteins provided by the mother in the yolk sac supports early development of the externally fertilized egg (27). Previous studies on *pomt1* and multiple transcriptomic studies defining maternal vs. zygotic mRNA expression showed that *pomt1* mRNA is highly expressed in the maternal deposit (20,36,37). In addition, work in the mouse found that glycosylated α-DG protein has a half-life of 25 days in muscle tissue (38). We hypothesized that the mRNA provided by the Het mother could provide sufficient maternal Pomt1 protein to glycosylate α-DG masking the appearance of dystroglycanopathy phenotypes in *pomt1* KO^Het^ fish.

Molecular investigation by Western blot at 5 dpf indicated that *pomt1* KO^Het^ larvae retained both the Pomt1 protein and substantial α-DG glycosylation (**Fig. 5A**). All survival studies on fish from Het X Het crosses were performed at a standard stocking density of 5 fish per liter and we wondered if reducing stocking density and competition for food would support better survival for the smaller KO^Het^ fish. We reduced to stocking density to around 3 fish per liter where no adverse effects of low-density housing were observed (39). While KO^Het^ still die by 1 year of age, we were able to obtain breeding age KO^Het^ fish and found that both KO^Het^ males and females were fertile. This allowed us to investigate whether offspring from oocytes lacking *pomt1* obtained from KO^Het^ female X Het male crosses (KO^KO^) would have more severe phenotypes. Note that this breeding scheme only yields heterozygous controls that will be designated as Het^KO^s to differentiate them from standard heterozygous (Het) fish. 5 dpf *pomt1* KO^KO^ larvae showed no residual Pomt1 expression (**Fig. 5B**). Since the zebrafish is known to compensate for genetic defects by upregulating related genes (40), we also analyzed expression of *pomt2* that acts in a complex with Pomt1 to initiate O-mannosylation (41). *pomt2* expression via qPCR showed high variability in Het fish from Het X Het crosses at 5 dpf and an increase in KO^Het^ larvae at 10 dpf similar to the increase in *pomt1* shown in **Fig. 1C** (**Suppl. Fig. 5A**). In the *pomt1* KO^KO^ larvae, *pomt1* mRNA expression was increased at 5 dpf, but decreased by 10 dpf and *pomt2* expression was unchanged at both timepoints (**Suppl. Fig. 5B**). However, despite the increase in *pomt1* mRNA observed in 5 dpf KO^KO^ larvae, Pomt1 was not present in protein lysates and α-DG glycosylation was absent at 5 dpf showing that the function of both proteins was completely lost (**Fig. 5B**). This result showed that the residual Pomt1 protein found in KO^Het^ larvae had indeed originated from the mother and that this contribution could be removed in KO^KO^ larvae.

**Figure 5.**
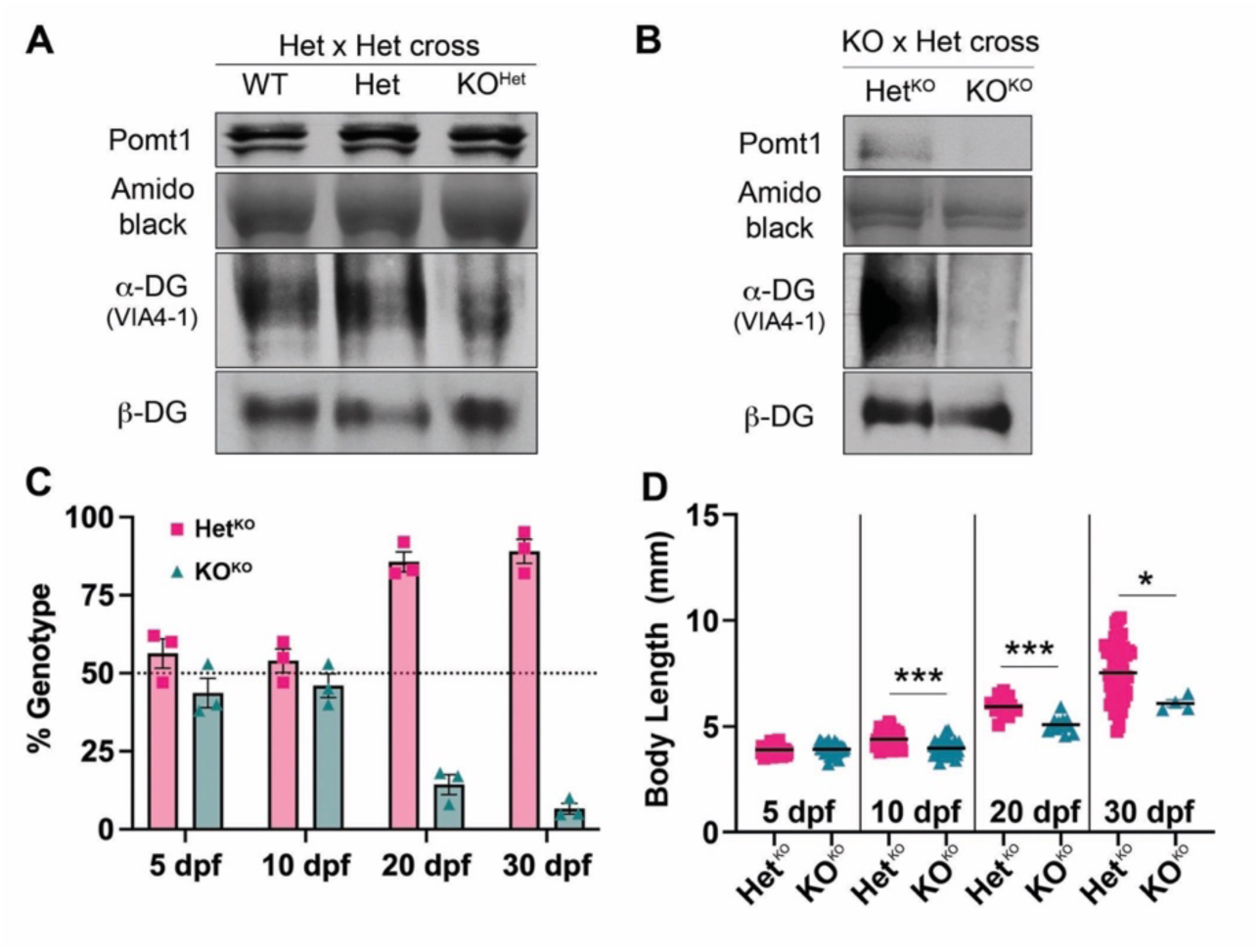
Removal of maternal *pomt1* mRNA contribution from oocytes leads to earlier phenotypes in *pomt1* KO^KO^ larvae. **A.** Residual Pomt1 is present in 5 dpf KO^Het^ larvae derived from Het X Het crosses, as is α-DG glycosylation. **B.** Both Pomt1 and α-DG glycosylation are absent at 5 dpf in KO^KO^ larvae obtained from KO mothers crossed with Het fathers. **C.** Survival in *pomt1* KO^KO^ zebrafish is reduced starting at 10 dpf as shown by the deviation from expected Mendelian ratios (50% for Het^KO^s and KO^Het^s) in 3 independent clutches. **D.** Reduced body size is observed in KO^KO^ larvae and juveniles starting at 10 dpf. *** p<0.001, * p<0.05

As observed in the *dag1* and *fkrp* mutant zebrafish (21,22), survival in *pomt1* KO^KO^ larvae decreased rapidly after 10 dpf (5 dpf: Het^KO^ 56.1±4.6%, n=167; KO^KO^ 43.9±4.6%, n=149; 10 dpf: Het^KO^ 54.0±3.8%, n=51, KO^KO^ 46.0±3.8%, n=43; 20 dpf: Het^KO^ 85.7±3.2% n=63, KO^KO^ 14.3±3.2% n= 12; 30 dpf: Het^KO^ 89.1±3.8% n=58, KO^KO^ 6.6±1.7% n= 8; N=3 clutches) (**Fig. 5C**). In parallel, body size was significantly reduced starting at 10 dpf (Het^KO^: 4.39±0.05 mm, n=51; KO^KO^: 3.97±0.05 mm, n=43, N=3 clutches. p<0.0001 ***) and was substantially smaller by 30 dpf in the few surviving animals (Het^KO^: 7.17±0.14 mm, n=58; KO^KO^: 6.22±0.20 mm, n=8, N=3 clutches. p<0.0001 ***) (**Fig. 5D**).

Locomotor activity was significantly impacted in *pomt1* KO^KO^ larvae at 5 dpf with reductions in distance traveled, velocity, and acceleration, showing that even if survival and size were not affected mobility was already severely impacted at an early age (Het^KO^ n=57, KO^KO^ n=36, N=3 clutches. Distance: He^KO^t: 345.2±117.3 cm, KO^KO^: 231.4±0.11, p<0.0001 ***; Velocity: Het^KO^: 0.19±0.06 cm/s, KO^KO^: 0.13±0.04 cm/s, p<0.0001 ***; Acceleration: Het^KO^: 166.6±67.6 cm/s^2^, KO^KO^: 81.63±67.34 cm/s^2^, p<0.0001 ***) (**Fig. 6A-C**). Disorganization of the muscle fibers and discontinuities in the basement membrane at myotendinous junctions (MTJ) between muscle fibers are major features of muscle disease caused by loss of binding between laminin and α-DG (17,21,42). We performed immunohistochemical staining with fluorescently stained phalloidin to outline actin in muscle fibers and laminin antibodies to label the basement membrane at MTJs in muscle cryosections from 5 and 10 dpf larvae. While there were no major differences at 5 dpf (**Suppl. Fig. 6A**), signs of severe muscle disease were found at 10 dpf with discontinuities in phalloidin staining and loss of laminin staining at the MTJs (**Fig. 6D**). Muscle fiber length was similar between *pomt1* Het^KO^ and KO^KO^ larvae at 5 dpf (**Fig. 6E**), but was significantly reduced at 10 dpf, even when length was normalized by total body length (Length: Het^KO^ 117.3±4.3 μm n=10, KO^KO^ 91.5±5.7 μm n=7, p=0.0023 **; Normalized length: Het^KO^ 1.00±0.04, KO^KO^ 0.86±0.05, p=0.045 *) (**Fig. 6F-G**). Measurement of fiber area was not feasible at 10 dpf because muscle fibers were too disorganized to obtain axial cross sections (**Suppl. Fig. 6B**).

**Figure 6.**
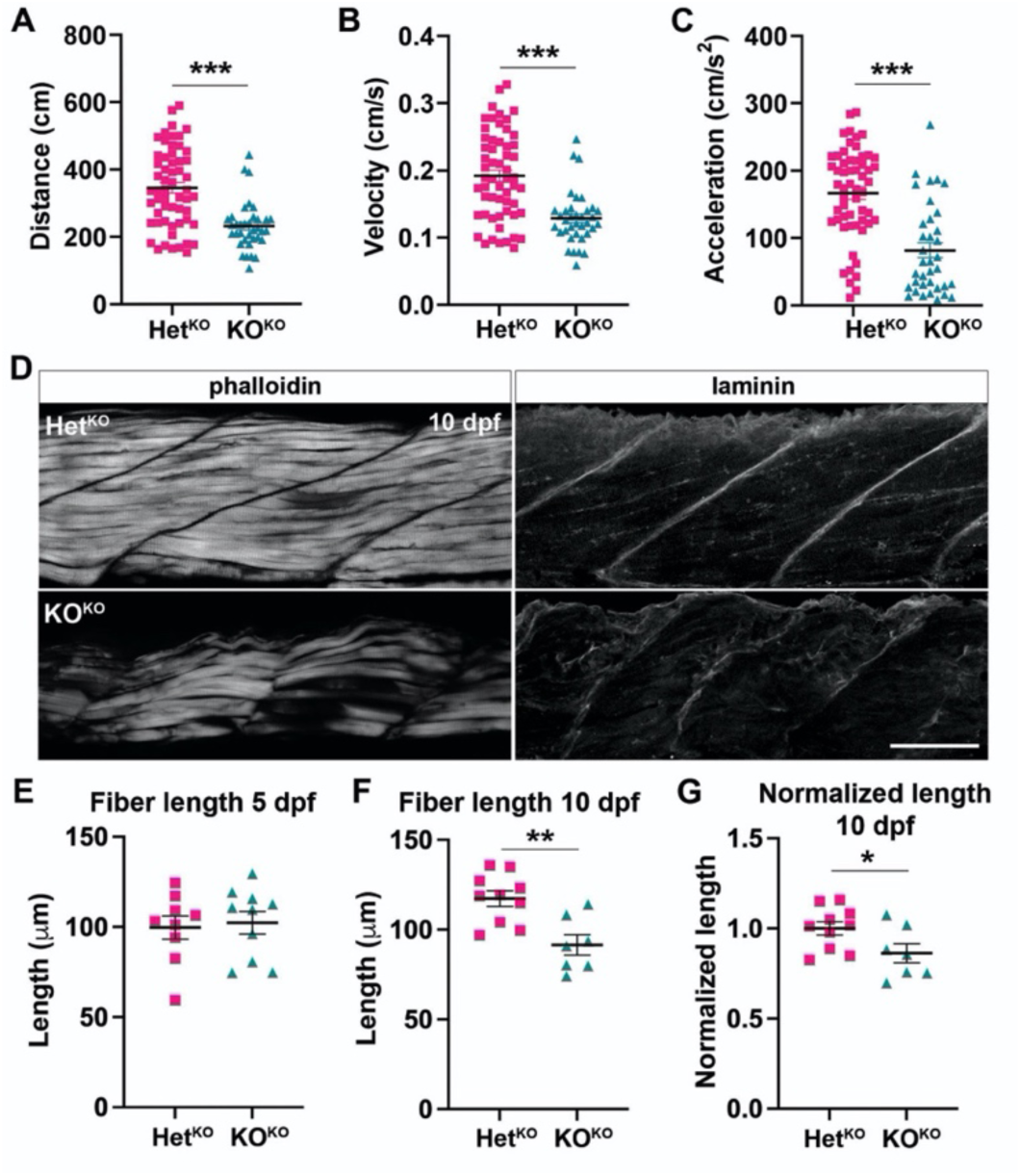
*pomt1* KO^KO^ larvae show reduced mobility at 5 dpf and muscle disease at 10 dpf. **A-C.** A significant reduction in total distance traveled (**A**.), velocity (**B.**) and acceleration (**C.**) are found in *pomt1* KO^KO^ larvae upon automated mobility analysis when compared to Het^KO^ siblings. *** p<0.001. **D.** Staining with fluorescently labeled phalloidin to visualize actin in muscle fibers revealed fiber disorganization from in the 10 dpf KO^KO^ larvae. Laminin staining was used to label the MTJ and showed discontinuities and disruption of the basement membrane between myomeres in KO^KO^s. See **Suppl. Fig. 6A** for 5 dpf muscle. Scale bar: 50 µm. **E-G.** Fiber length was the same at 5 dpf (**E.**), but it was significantly reduced at 10 dpf (**F.**) even when length was normalized for body length (**G.**). ** p<0.01, * p<0.05

We conducted further analysis of muscle ultrastructure at 10 dpf using transmission electron microscopy (TEM). As fibers in the same muscle can be differentially affected, multiple fibers were analyzed per animal. Results were consistent with other CMD zebrafish mutants (21,43,44), showing consistent swelling of the SR with misalignment of adjacent sarcomeres in the most severe cases (**Fig. 7A**). Triad area was increased (Het^KO^ 13.01±0.63 nm^2^ n=17 fibers/387 triads, KO^KO^ 15.54±0.74 nm^2^ n=17 fibers/435 triads, p=0.0076 **; N=4 animals) (**Fig. 7D**). To represent variability, we scored images as organized (aligned sarcomeres) or disorganized (misaligned sarcomeres of variable size) and quantified the percentage of disorganization per animal (**Fig. 7E**). Triad area was significantly increased in both organized and disorganized KO^KO^ fibers (**Suppl. Fig. 6C**). Sarcomere length was reduced (Het^KO^ 2.08±0.03 μm n=19 fibers/349 sarcomeres, KO^KO^ 1.93±0.03 μm n=18 fibers/333 sarcomeres, p=0.0009 ***; N=4 animals) (**Fig. 7F**). Separation of adjacent myofibers was observed in the KO^KO^ muscle with swelling and rupture of mitochondria (**Fig. 7B**). The MTJ showed lower electron density due to the likely loss of collagen and ECM as also shown in the laminin staining in **Fig. 6D** and discontinuities at the border with myofibers (**Fig. 7C**).

**Figure 7.**
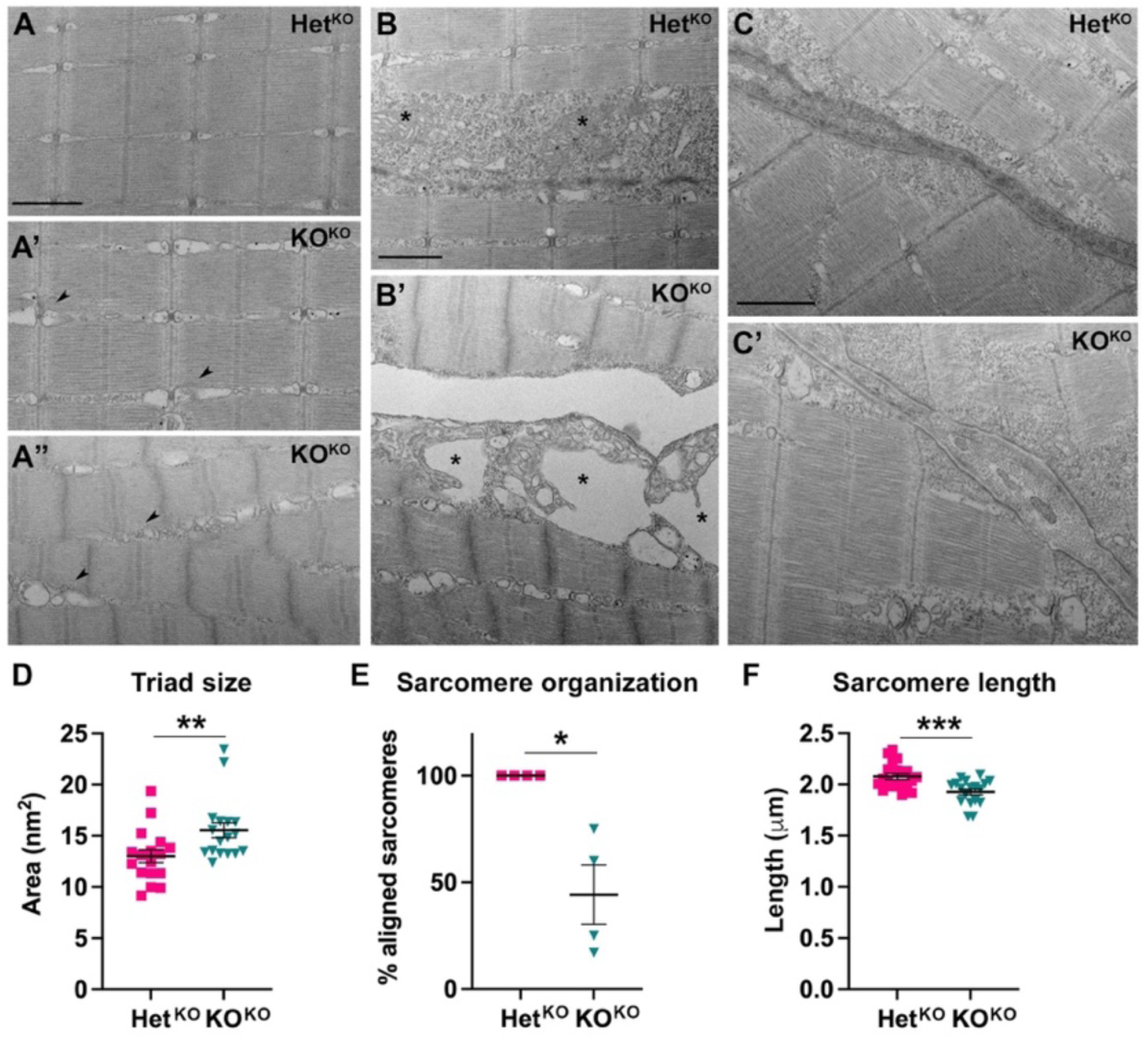
TEM analysis of *pomt1* KO^KO^ muscle at 10 dpf. **A.** Sarcomeres showed variable severity of disruptions in KO^KO^ larvae with enlargement of the SR (arrowheads in **A’** and **A”**) and misalignment of the sarcomeres in the most severely affected fish (**A”**). **B.** The boundary between myofibers was detached in the KO^KO^ (**B’**) with severe swelling and membrane disruption of the mitochondria. For comparison mitochondria are labeled with asterisks in both the Het^KO^ (**B**) and KO^KO^ (**B’**). **C.** The MTJ showed reduced electron density and discontinuities in the border where myofibers are attached (white arrows in **C’**). Scale bars: 500nm. **D.** Quantification of triad area in independent muscle fibers from 4 larvae per genotypes showed a significant increase. See **Suppl. Fig. 6C** for quantification in moderately (**A’**) and severely (**A”**) affected fibers. **E.** Quantification of the percentage of fibers with aligned (**A’**) or misaligned (**A”**) sarcomeres showed variability as expected for muscle disease, but all fish were affected. **F.** Sarcomere length was significantly reduced in the *pomt1* KO^KO^ larvae. *** p<0.001, ** p<0.01, * p<0.05

Finally, we analyzed the neuromuscular junction (NMJ) in 10 dpf muscle using whole mount staining with α-bungarotoxin (α-BTX) to label postsynaptic acetylcholine receptors (AChRs) in the motor endplate and immunostaining using an anti-SV2 antibody to visualize presynaptic terminals. α-BTX staining showed an overall reduction in intensity in the muscle fibers in the myomere and at MTJs in *pomt1* KO^KO^ larvae (Intensity myomere: Het^KO^ 0.30±0.02 n=11 larvae, KO^KO^ 0.20±0.02 n=13 larvae, p=0.0021 **. Intensity MTJ: Het^KO^ 0.34±0.02 n=11 larvae, KO^KO^ 0.24±0.02 n=13 larvae, p=0.0018 **) (**Fig. 8A-B**). There was also an increase in α-BTX puncta number in the myomere suggesting NMJ fragmentation, while puncta density at the MTJ did not show significant differences (α-BTX puncta/100 μm^2^ myomere: Het^KO^ 3.20±0.25 n=11 larvae, KO^KO^ 3.55±0.21 n=13 larvae, p=0.013 *. α-BTX puncta/100 μm^2^ MTJ: Het^KO^ 0.34±0.02 n=11 larvae, KO^KO^ 0.24±0.02 n=13 larvae, p=0.30) (**Fig. 8C**). There was an overall decline in SV2 puncta (SV2 puncta/100 μm^2^ myomere: Het^KO^ 1.03±0.08 n=11 larvae, KO^KO^ 0.43±0.10 n=13 larvae, p=0.0004 ***. SV2 puncta/100 μm^2^ MTJ: Het^KO^ 2.57±0.21 n=11 larvae, KO^KO^ 1.68±0.29 n=13 larvae, p=0.03 *) (**Fig. 8D**). As expected, the colocalization coefficient for α-BTX and SV2 was also significantly reduced (Colocalization myomere: Het^KO^ 0.61±0.04 n=11 larvae, KO^KO^ 0.34±0.05 n=13 larvae, p=0.0007 ***. Colocalization MTJ: Het^KO^ 0.70±0.02 n=11 larvae, KO^KO^ 0.41±0.05 n=13 larvae, <p=0.0001 ***) (**Fig. 8E**). Overall, the 10 dpf KO^KO^ muscle presented with multiple features consistent with severe muscle disease.

**Figure 8.**
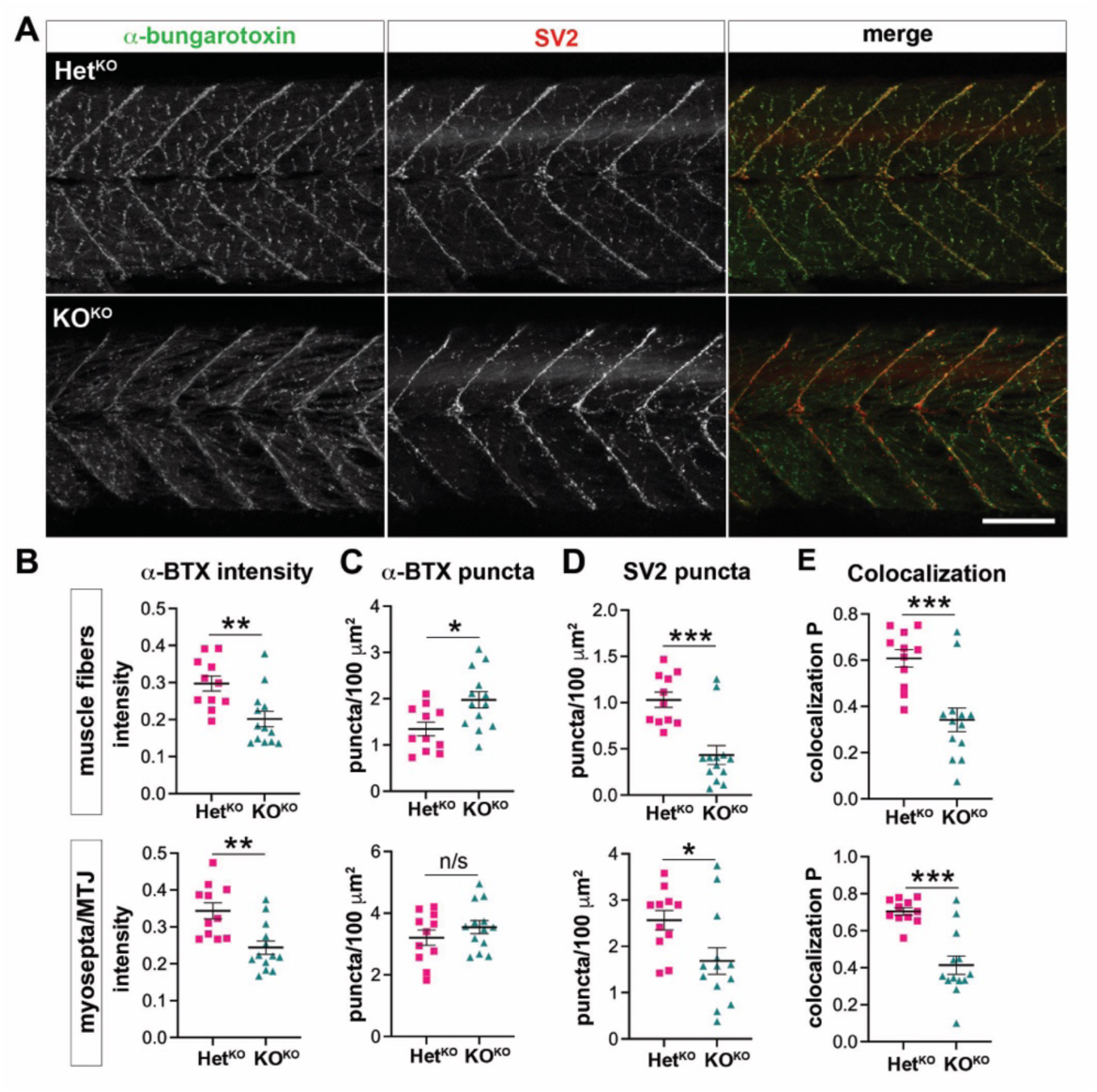
Neuromuscular junction (NMJ) analysis in 10 dpf *pomt1* KO^KO^ larvae. **A.** Whole mount staining with α-bungarotoxin (α-BTX, green) for acetylcholine receptors and SV2 for presynaptic sites showed both changes in intensity and organization in KO^KO^ larvae. Scale bar: 100 µm. **B.** α-BTX staining intensity was reduced both on muscle fibers in the myomeres and at the myosepta. **C**. α-BTX puncta number was increased on the muscle fibers. **D.** SV2 puncta were reduced at both sites. **E.** The Pearson coefficient of colocalization between the two markers was significantly reduced in both. *** p<0.001, ** p<0.01, * p<0.05

### Retinal synapse formation and axon guidance deficits are found in pomt1 KO^KO^s at 5 dpf

The same retinal phenotypes observed at 30 dpf in KO^Het^ fish were present at 5 dpf in the KO^KO^ larvae. Arr3a staining in cone photoreceptors was patchy and nuclei were disorganized in the outer nuclear layer of photoreceptors (**Fig. 9A**). Synaptic staining with Syp in the OPL was discontinuous, often extending into the outer nuclear layer indicating retraction of photoreceptor pedicles indicating an early loss of synapses between photoreceptors and bipolar cells (**Fig. 9A**, **Suppl. Fig. 7A**). In addition, discontinuities in horizontal cell dendrite distribution in the OPL were noted when these cells were labeled using an anti-calbindin antibody (**Fig. 9B**).

**Figure 9.**
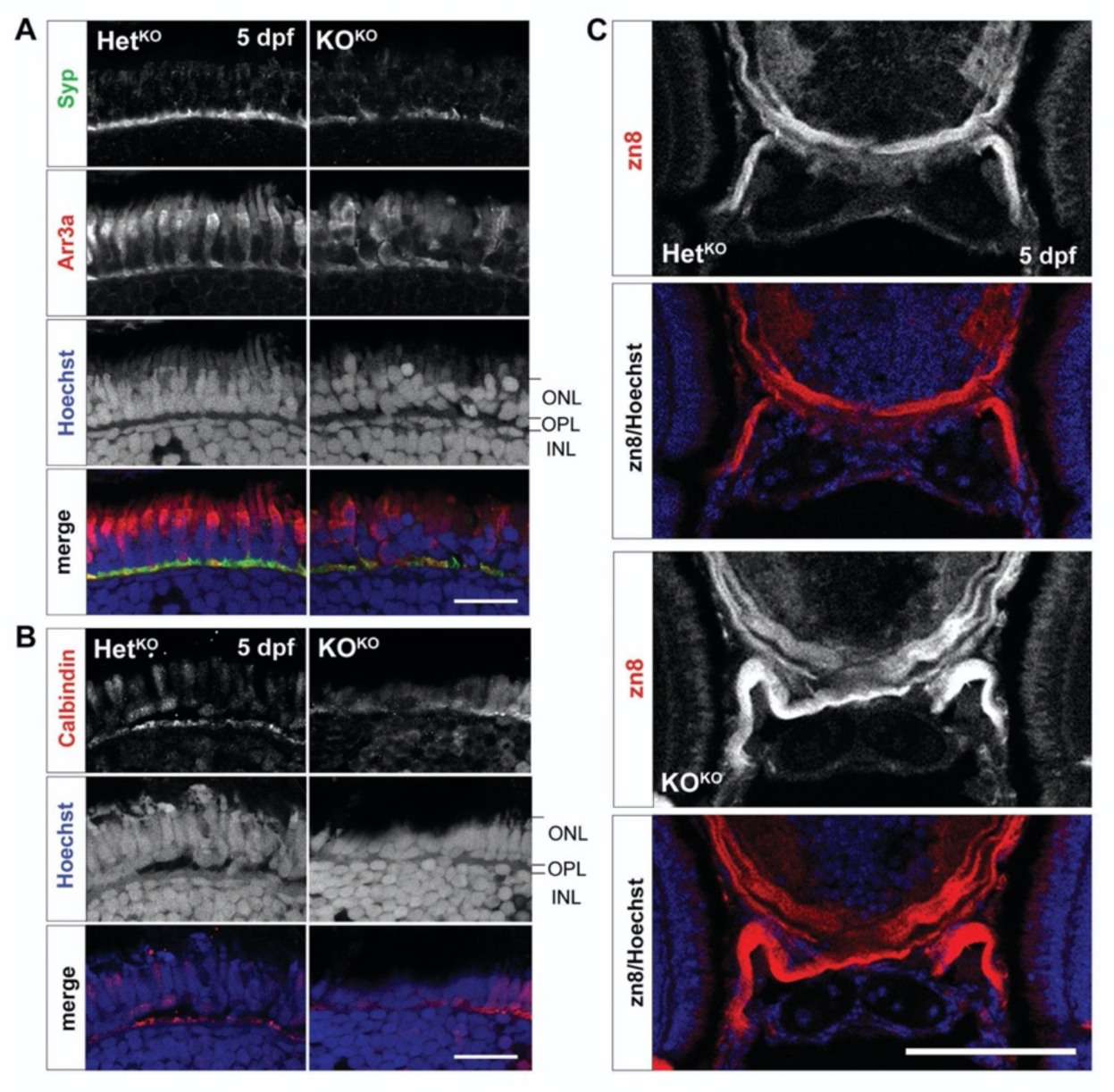
Retinal and axon guidance phenotypes are found at 5 dpf in *pomt1* KO^KO^ larvae. **A.** Discontinuities in synaptophysin (Syp) staining are found in the outer plexiform layer (OPL) of *pomt1* KO^KO^ retinae. Red/green double cone marker arrestin 3a (Arr3a) again shows disorganized outer segments and patchy staining and altered shape of photoreceptors pedicles. Nuclear staining with Hoechst shows disorganization disruption in the photoreceptor nuclei in the outer nuclear layer (ONL) and in horizontal cell nuclei at the top of the inner nuclear layer (INL). Scale bar: 20 µm. **B.** Horizontal cell dendrites extending in the OPL are visualized using anti-calbindin staining and are disrupted in the KO^KO^ retina. Scale bar: 20 µm **C.** The optic nerve was labeled using the zn8 antibody that marks retinal ganglion cell axons showing axon defasciculation after optic chiasm crossing. Imaged including the full head are shown in **Suppl. Fig. 8B.** Scale bar: 50 µm.

We then explored additional developmental phenotypes. Deficits in the inner limiting membrane (ILM), a basement membrane lining the retinal ganglion cell (RGC) layer, which is involved in RGC polarization and migration, were noted in dystroglycanopathy mouse models (45). The developing zebrafish ILM is very thin due to the direct opposition of the lens to the RGC layer and *pomt1* KO^KO^ larvae showed no differences in laminin fluorescent signal and appeared indistinguishable from Het^KO^ siblings in respect to the inner retina (**Suppl. Fig. 7B**). One major phenotype caused by loss of α-DG glycosylation in mice is the disruption of axonal tracts in the brain caused by loss of interaction with axon guidance cues (46,47). One example is the disruption in RGC axon sorting and fasciculation in the optic nerve which is controlled by binding to the midline crossing cue Slit2, an α-DG ligand (47,48). We did not see any deficits in axonal crossing and fasciculation in the optic chiasm of 30 dpf KO^Het^ fish (**Suppl. Fig. 8A**), but we wondered whether the maternal deposit of *pomt1* was sufficient to bypass this early developmental phenotype. When we analyzed the optic chiasm using the zn8 antibody that labels retinal ganglion cells and their axons, we noted substantial axonal defasciculation following axon crossing at the optic chiasm of KO^KO^ larvae reminiscent of phenotypes found following *slit2* knockout and knockdown in the fish (**Fig. 9C, Suppl. Fig. 8B**) (49). Thus, *pomt1* loss of function early in development in the zebrafish also recapitulates some of the central nervous system phenotypes found in higher vertebrates.

## Discussion

Here, we showed that *pomt1* KO zebrafish can be used to model dystroglycanopathy phenotypes in the muscle, eye, and brain with variable severity. For our initial analysis of this mutant strain, we focused on phenotypes identified in other dystroglycanopathy models in fish and mouse. The phenotypes found in *pomt1* KO^Het^ obtained from heterozygous crosses were milder than those observed in the *dag1* loss of function fish (21) suggesting that compensation may be present. We found that this was due to high *pomt1* mRNA expression in the maternal deposit in the oocyte supporting α-DG glycosylation during the first week post fertilization. Due to external fertilization, the early embryo has been shown to exclusively rely on maternally deposited RNA transcripts and proteins until the maternal-zygotic transition between 2.7 and 3.7 hours post-fertilization (hpf) or 512-cell to oblong stage (50). Avsar-Ban et al. have shown that the mRNA level of *pomt1* is five-fold higher at 0 hpf (prior to zygotic generation) than at 6 hpf (zygotic generation) (20). Additional transcriptional analyses to determine maternal vs. zygotic contribution have also identified *pomt1* as a maternally provided transcript (36,37). In mouse tissue glycosylated dystroglycan protein has a half-life of 25 days (38), a similar half-life in the zebrafish would overlap zygotic transition and through early development. While we still found Pomt1 expression at 5 dpf in KO^Het^ larvae, the protein was largely absent from the muscle at 10 dpf. In parallel, *pomt1* mRNA levels that were consistently lower than WT at all other timepoints were increased at 10 dpf likely in response to loss of the maternal protein with a corresponding increase in *pomt2*. We found that in *pomt1* KO^Het^ fish α-DG glycosylation was completely lost by 30 dpf when disease-related phenotypes in the muscle and retina began to emerge. KO^Het^ juveniles maintained at high stocking density started dying at 30 dpf with no survivors found past 70 days. Careful and continuous size segregation and reduced stocking density allowed survival of KO^Het^ mutants to breeding age. These results indicate that even when the early developmental roles of α-DG are conserved, continued Pomt1 expression is necessary to maintain retinal photoreceptors integrity. This is consistent with findings in *pomt2* and *pomgnt1* mutants, that show similar delayed phenotypes (23,24) and are likely to undergo the same maternal compensation due to high levels of zygotic mRNA identified in transcriptomic studies (36,37).

By raising KO females, we were able to obtain oocytes completely devoid of *pomt1* mRNA to generate true developmental KOs where α-DG O-mannosylation is prevented from time of fertilization. *pomt1* KO^KO^s closely resembled other dystroglycanopathy models in both zebrafish and mouse. As in the *dag1* mutant, *patchytail*, *pomt1* KO^KO^s died within the first 2 weeks post fertilization. While these larvae showed substantial mobility phenotypes at 5 dpf, muscle integrity was preserved which is consistent with the progressive loss of muscle integrity shown in *dag1* mutants by birefringence assays (21). Multiple deficits in muscle organization and ultrastructure were evident at 10 dpf when the fish begin to die, including loss of MTJ integrity, enlarged SR triads, and disorganized sarcomeres. These features are consistent with ultrastructural analysis of muscle obtained from mouse models deficient for *Dag1* and individuals with dystroglycanopathy (51,52). Comprehensive NMJ analysis had only been performed on morphant larve with *fkrp* knockdown also showing extensive fragmentation of AChR clusters (53). In addition, we identified a loss of SV2 staining suggesting a concurrent reduction in pre-synaptic terminals. The first α-DG ligand ever identified was Agrin, a major organizer of the NMJ, and this interaction was shown to regulate AChR aggregation and stability (54,55).

As observed in *pomt1* KO^Het^ fish, photoreceptor synapse loss is present earlier at 5 dpf in KO^KO^ larvae in conjunction with defects in optic tract fasciculation reflecting loss of α-DG ligand binding necessary for synapse formation and axon guidance that had been previously described in *pomt1* conditional mouse models and *pomgnt1* mutant zebrafish (13,23). α-DG can bind to a variety of proteins containing laminin-G domains (56) and several of its ligands have important roles in axon guidance and synapse formation in the brain and peripheral nervous system (46,48,57). Slits are involved in guiding axons in crossing the midline of the body and contribute to the formation of the optic chiasm where axons from each eye target brain regions on the opposite side (58). α-DG was shown to bind to the laminin-G domain of Slit2 to organize axonal crossing (48) and the phenotypes observed in the *pomt1* KO^KO^ larval optic chiasm are similar to the enlarged and disorganized axonal tract found in *slit2* KO zebrafish (59). Another α-DG ligand is pikachurin, a postsynaptic adhesion protein involved in the formation and maintenance of photoreceptor ribbon synapses (57,60). Ribbon synapses are highly specialized structures in the photoreceptor pedicles in the OPL mediating rapid transmission of visual signals to the dendrite tips of horizontal cells and bipolar cells. When the bond between α-DG and pikachurin is lost, the synapse comes apart and photoreceptors pedicles retract eventually leading to photoreceptor degeneration and loss of vision (13,57,60). Our findings in *pomt1* KO^KO^s closely recapitulate phenotypes in a mutant mouse line where *Pomt1* is conditionally removed in photoreceptors (13). In addition, α-DG had been shown to interact with the secreted ECM protein eye shut homolog (EYS) which is also involved in photoreceptor survival and is frequently mutated in retinitis pigmentosa (23,61). We observed no major disruptions in retinal laminar organization, but additional analyses will be needed to determine whether synapses in other retinal layers are also affected as described in mouse models (45). No deficits were observed in the inner limiting membrane lining the RGC layer, but this finding may be due to anatomical differences between the mouse and the zebrafish and further detailed analyses on other synapses and RGCs are needed. Overall, early developmental phenotypes caused by loss of matriglycan binding in multiple tissues are present in the *pomt1* KO^KO^ supporting the need to further study of this model to determine the impact of these mutations in the brain and peripheral nervous system.

While *pomt1* KO^KO^ and the *dag1* ENU mutant are similar, their phenotypes are overall less severe than those observed the *fkrp* mutant line which in turn recapitulate morphant models for *fkrp* and other glycosyltransferases mutated in dystroglycanopathies (17,18,22,62). Morphants for *fkrp*, *pomgnt1* and *b3galnt2* larvae show early mortality, shorter body, smaller head and eyes and severe cardiac edema. On the other hand, *pomt1* morphants were less severe than other morphants showing milder morphological disruptions (20). Thus, we believe that KO^KO^ animals reflect complete loss of function of *pomt1* in the fish and can be studied as a valid model of dystroglycanopathy phenotypes in the muscle, eye, and brain. Furthermore, given the spectrum of disease in *POMT1* patients compared to dystroglycanopathies caused by alternative genes, we hope to leverage our model to identify potential novel cellular deficits that can inform pathological analysis in individuals with *POMT1* mutations and identify novel targets for therapeutic intervention. Our study stresses how maternal mRNA and protein contribution must be taken into consideration when generating dystroglycanopathy models since the long-lasting presence of glycosylated α-DG can mask developmental phenotypes in the embryo. Similarly mild phenotypes identified in *pomgnt1* and *pomt2* zebrafish KO obtained from heterozygous crosses (23, 24) could also be due to maternal compensation since both these genes are also maternally provided (36, 37). At the same time, these later onset models where pathogenesis progresses more slowly like the *pomt1* KO^Het^ fish could still be used to study the involvement of α-DG glycosylation in maintenance of tissue function and to investigate therapeutic interventions in multiple tissues at different stages and severity of disease.

## Materials and Methods

### Zebrafish Husbandry

All experiments and procedures were conducted according to institutional guidance and approved by the Rutgers University Institutional Animal Care and Use Committee. The *pomt1* mutant line animals were generated from the Zebrafish Mutation Project (line: sa146678, ZFIN ID: ZDB-ALT-130411-3355) on an AB background (28) The line was a kind gift of Dr. James Dowling (Sick Kids, Toronto, Canada) and it was received after 2 subsequent AB outcrossings to ensure genetic cleanup of secondary off-target mutations. The breeding stocks were then bred for four generations prior to experimental use while housed in recirculating Tecniplast USA systems under a 14/10 h light/dark (LD) cycle at 28°C and fed twice daily. To produce embryos, male and female zebrafish were paired in the evening, and spawning occurred within 1 h of lights-on the following morning. Embryos were placed in 10 cm petri dishes with egg water containing methylene blue (5 x 10^-5^ % w/v) and raised under LD cycles at 28°C. Initial stocking densities included 50 larvae, 25 juveniles and 15 adults per 3.5 liter tanks. In order to raise and maintain KO mutants, careful and continuous size segregation and reduced stocking density were conducted (25 larvae, 15 juveniles and 8 adults). Only clutches in which >10% of the wild-type or heterozygous siblings were non-viable were utilized for experimentation.

### Genotyping

DNA from larval tails and juvenile and adult tail fins was extracted using the Extract-N-Amp Tissue PCR Kit (Millipore Sigma). After verifying SNP location with Sanger sequencing at Azenta Life Sciences, genotype was determined using a Custom TaqMan SNP Genotyping Assay (Assay ID: ANRWEV2, Thermo Fisher).

### Quantitative Real-Time PCR (qPCR) Gene Expression Analysis

Total RNA was isolated from embryo composite samples or adult tissue using ReliaPrep RNA Tissue Miniprep System (Promega). Reverse transcription was performed on RNA to produce cDNA for RT-PCR using an iScript Reverse Transcription Supermix (Bio-Rad) using a Bio-Rad T100 Thermal Cycler. RT-PCR reactions were performed in triplicate using PowerUp SYBR Green Master Mix (Thermo Fisher). cDNA amplification was performed for 40 cycles on a QuantStudio 3 (Applied Biosystems) and recorded with QuantStudio Design and Analysis Software. Analysis was conducted as ΔΔCT using *elf1-alpha* or *rpl13* as an endogenous control. All primer sequences are available upon request.

### Wheat Germ Agglutinin Enrichment and Western Blot Analysis

Total zebrafish protein was extracted from composite samples (5 dpf; N=60 and 30 dpf; N=3) using lysis buffer (50 mM Tris pH 8, 100 mM NaCl, 1 mM PMSF, 1 mM Na Orthovanadate, 1% Triton-X 100) containing protease inhibitor cocktail (Millipore Sigma), and 3 rounds of hand homogenization and vortexing. Samples were then sonicated in a Bransonic B200 Ultrasonic Cleaner (Millipore Sigma) 3 times for 5 minutes. Next samples were spun down at 4°C for 20 minutes at 15,000 xg. The supernatant was collected from the samples and sonicated once for 5 minutes. Total protein concentration was determined using a BCA Protein Assay kit (G-Biosciences) and subsequently treated with DNase I (New England Biolabs). 400-500 ±g of protein was then added to 50 µl of Wheat Germ Agglutinin (WGA) agarose bound beads (Vector Laboratories) filled to a volume of 300 µl with Lectin Binding Buffer (20 mM Tris pH 8, 1 mM MnCl_2_ and 1 mM CaCl_2_) in a 1.5 ml tube and rocked overnight at 4°C. Samples were centrifuged at 15,000 xg for 2 minutes and the supernatant was collected. Volumes of supernatant containing 60 µg of protein were set aside for detection of Pomt1. Glycoproteins were eluted by boiling at 100°C for 10 minutes using 4x Laemmli sample buffer (Bio-Rad). Proteins were separated using a 4-12% Bis-Tris Protein SDS-PAGE Gel (Life Technologies). Samples were transferred to a nitrocellulose membrane (Thermo Fisher) and stained with 0.1% naphthol blue black (amido black) (Millipore Sigma) to detect protein transfer and for total protein quantification. The membranes were blocked for 1 hour with 5% milk in Tris-Buffered Saline with 0.1% Tween (TBST) and subsequently probed with 1:100 anti-α-Dystroglycan antibody, clone VIA4-1 (05-298, Millipore Sigma) and 1:1,000 anti-lll-Dystroglycan antibody (ab62373, Abcam) or anti-POMT1 (A10281, ABclonal) overnight at 4°C. Then probed with 1:3,000 peroxidase AffiniPure donkey anti-mouse IgG and 1:20,000 peroxidase AffiniPure donkey anti-rabbit IgG (Jackson ImmunoResearch). After washing with TBST, the blots were developed using chemiluminescence Pierce ECL Western Blotting Substrate on CL-XPosure™ Film (Thermo Fisher).

### Morphological Analysis

Larvae and juvenile zebrafish were anesthetized in 0.016% w/v tricaine methane sulfonate (Tricaine, MS-222) and immobilized in 5% methyl cellulose for imaging with a M165 FC stereo microscope and LAS Software V4.21 (Leica Microsystems). Total body length measurements and eye area were quantified using Fiji/ImageJ software (63). Retinal and photoreceptor layer thickness were measured on cryosections prepared as described below and stained with Hoechst 33342 (1:10,000, Thermo Fisher) also using Fiji/ImageJ.

### Automated Mobility Tracking

Automated mobility assays were conducted by randomizing 5, 20, and 30 dpf zebrafish into sterile 96 and 24-well plates filled with 200 µl and 2 ml of system water, respectively. Fish were then habituated in the DanioVision^TM^ Observation Chamber (Noldus Information Technology, Wageningen) for 30 minutes at a controlled temperature of 28°C. Locomotor activity was tracked for 30 minutes using EthoVision® XT video tracking software. A minimum of three unique breeding sets were used to compile results at each timepoint.

### Fluorescent Immunohistochemisty

Zebrafish larvae at 5 dpf were euthanized with tricaine and a small portion of the tail was cut to use for genotyping. The remaining fish body was fixed in 4% paraformaldehyde overnight at 4°C. Larvae were washed three times in 1x PBS, then cryoprotected in 15% sucrose with 0.025% sodium azide followed by 30% sucrose with 0.025% sodium azide. Larvae were embedded in Tissue Freeze Medium (TFM) (General Data Healthcare) and frozen in 2-methyl butane with dry ice. Juvenile fish processing required the addition of a decalcification step in Cal-Ex (Fisher Scientific) following fixation for 1 hour and 15 minutes rocking at 4°C. Cryosections were performed using a Leica CM1850 UV Cryostat (Leica Microsystems) at 10 μm for 5 dpf, 12 μm for 10 dpf, and 16 μm for 30 dpf. Primary antibodies were applied overnight at 4°C in 1% normal goat serum (NGS) with 0.1% Triton following a blocking step with 10% NGS. Secondary immunostaining was applied for 1 hour. The following antibodies were used: primary -POMT1 (A10281, 1:100, ABclonal), Gnat2 (A10352, 1:200, ABclonal), Synaptophysin (ab32127, 1:100, Abcam), Ryr1 (MA3-925, 1:100, Thermo Fisher), zpr-1 (Arr3a) (zpr-1, 1:200, ZIRC), zpr-3 (zpr-3, 1:200, ZIRC), zn-8 (zn-8-s, 1:100, DSHB), Laminin (L9393, Sigma); secondary - goat anti-mouse IgG Alexa Fluor 568, goat anti-rabbit Alexa Fluor 488 or goat anti-rabbit Alexa Fluor 568 (all from Thermo Fisher). Slides were washed twice in 1x PBS at room temperature and Hoechst 33342 (1:10,000, Thermo Fisher) was applied in 1x PBS for 5 minutes. Staining of F-actin/phalloidin was performed using Alexa Fluor™ 546 Phalloidin (1:400; Invitrogen™) in 1% NGS with 2% Triton X and kept in the dark for 1 hour at room temperature. Slides were then washed three times in 1x PBS for 5 minutes and countered stained with DAPI (1:20,000, Thermo Fisher). Slides were coverslipped using ProLong™ Gold Antifade Mountant (Thermo Fisher). For whole mount staining fixed zebrafish were permeabilized in 1 mg/mL collagenase D (Sigma) for 1.5 hours at RT. Alexa Fluor™ 488 α-Bungarotoxin conjugate (Invitrogem) was diluted 1:500 in antibody block (5% BSA, 1% DMSO, 1% Triton, 0.2% saponin in 1x PBS) for 2 hours at RT following several washes of PBS-0.1% Tween. Samples were then blocked overnight at 4°C before being incubated in 1:50 SV2 antibody (DHSB) in block at RT for 8 hrs then for 48 hours at 4°C 4. The samples were rinsed with PBS-0.1% tween followed by 1:250 dilution of secondary staining overnight at 4°C. The secondary antibody was rinsed with PBS-0.1% tween before mounting in 1.5% low melt agarose at 80°C for imaging. Images were for histological and whole-mount staining were taken using a Zeiss LSM800 confocal microscope and Zeiss Zen imaging software.

### Paraffin Sectioning and Hematoxylin & Eosin Staining

Paraffin sectioning was performed by the Rutgers Cancer Institute of New Jersey Biospecimen Repository and Histopathology Service Shared Resource. Decalcified juvenile fish were placed in 70% Ethanol. Specimens were then dehydrated through a step protocol using ethanol and xylene and embedded in paraffin using a Sakura Tissue-Tek VIP6 AI tissue processor and a Sakura Tissue-Tek TEC 6 tissue embedder. Each mold was sectioned on a Leica Reichert-Jung BioCut 2030 rotary microtome at 4 µm. Slides were then deparaffinized and stained with hematoxylin and eosin using a Sakura Tissue-Tek DRS 2000 slide stainer.

### Transmission Electron Microscopy (TEM)

Zebrafish larval were fixed 4% Paraformaldehyde, 2.5% Glutaraldehyde, 0.1M Sodium Cacodylate Buffer, and 8mM CaCl_2_ (All from Electron Microscopy Sciences) overnight at 4C. Post-fixation and embedding were conducted by New York University (NYU) Langone’s Microscopy Laboratory. Tissue obtained by NYU was rinsed in 0.1 M Cacodylate buffer and post-fixed in 1% Osmium tetroxide in 0.1M Cacodylate buffer. Samples were rinsed again in ddH2O and dehydrated in a series of ethanol washes. Following dehydration, they were then incubated in two series of EMbed 812 with the addition of N-Benzyl-N, N-Dimethylamine (BDMA) after initial incubation. Samples were embedded and cut longitudinally 95 nm thickness thin section on 100 nm formvar-coated Cu Grids and stained with 0.2% lead citrate. Sections were then imaged at the Rutgers Electron Microscopy Core Imaging Laboratory Philips CM12 electron microscope with AMT-XR11 digital camera.

### Image Quantification and Statistical Analysis

All imaging analysis was performed with the investigator masked to the genotype on randomized images. Quantification of myotome length was performed by measuring distance between myosepta on H&E or fluorescent images for at least 8 myosepta per fish using Fiji/ImageJ. Analysis of α-BTX and SV2 staining was conducted through a custom CellProfiler (64) pipeline collecting puncta intensity, number, and size information on at least 4 myomeres and myosepta per fish. TEM images were analyzed by manually tracing 18-20 triads and sarcomeres per image using Fiji/ImageJ on 4-5 independent muscle fibers per fish. Data represent the mean ± SEM or are presented as scatter plots with the mean. All statistical analyses were performed using GraphPad Prism v.8.20 (GraphPad, San Diego, CA). To compare *in vivo* data for two groups, individual means were compared using the MannWhitney U test and Welch’s t test. When comparing more than three groups, normality was assessed using the Shapiro-Wilk test, then a two-way ANOVA with Tukey’s post hoc test was used for normally distributed samples or a Kruskal-Wallis test with Dunn’s post-hoc test was used for non-normally distributed samples. All statistical analyses were two-sided tests, and p values of ≤ 0.05 were considered statistically significant.

## Acknowledgements

The authors would like to thank Dr. James Dowling (Sick Kids, Toronto, ON, Canada) for donating the sa146678 mutant line. We also thank Drs. John Dowling (Harvard University) and Ellen Townes-Anderson (New Jersey Medical School) for helpful discussion on zebrafish photoreceptor phenotypes and retinal degeneration, and Dr. Matthew Alexander (University of Alabama, Birmingham) and Dr. Clarissa Henry (University of Maine, Orono) for discussion on the muscle phenotypes. We are particularly grateful to Kathleen Flaherty at the Rutgers Zebrafish Facility with assistance in the maintenance of the zebrafish colony and Lucyann Franciosa and Kelly Watkinswalton at the Rutgers Cancer Institute of New Jersey Biospecimen Repository and Histopathology Service Shared Resource for paraffin sectioning. We would also like to thank Dr. Feng-Xia (Alice) Liang at New York University Langone Health’s Microscopy Laboratory for sample preparation for electron microscopy and Dr. Rajesh Patel of the Rutgers Core Imaging Laboratory for use of the TEM microscope. The zn-8 developed by University of Oregon, Department of Neuroscience was obtained from the Developmental Studies Hybridoma Bank, created by the NICHD of the NIH and maintained at The University of Iowa, Department of Biology, Iowa City, IA 52242.

## Abbreviations

α-DG: α-dystroglycan
CMD: congenital muscular dystrophy
dpf: day post fertilization
ECM: extracellular matrix
Het: heterozygous
KO: knock-out
LGMD: limb-girdle muscular dystrophy
MO: morpholino oligonucleotides
MTJ: myotendinous junction
NMJ: neuromuscular junction
OPL: outer plexiform layer
RGC: retinal ganglion cell
SR: sarcoplasmic reticulum
TEM: transmission electron microscopy
WT: wild type
WWS: Walker Warburg Syndrome

## References

1. Bönnemann, C.G., Wang, C.H., Quijano-Roy, S., Deconinck, N., Bertini, E., Ferreiro, A., Muntoni, F., Sewry, C., Béroud, C., Mathews, K.D., et al. (2014) Diagnostic approach to the congenital muscular dystrophies. Neuromuscular disorders : NMD, Vol. 24, pp. 289–311.

2. Kanagawa, M. (2021) Dystroglycanopathy: From Elucidation of Molecular and Pathological Mechanisms to Development of Treatment Methods. Int J Mol Sci, 22, 13162.

3. Yoshida-Moriguchi, T. and Campbell, K.P. (2015) Matriglycan: a novel polysaccharide that links dystroglycan to the basement membrane. Glycobiology, 25, 702–713.

4. Brancaccio, A. (2019) A molecular overview of the primary dystroglycanopathies. J Cell Mol Med, 23, 3058–3062.

5. Zambon, A.A. and Muntoni, F. (2021) Congenital muscular dystrophies: What is new? Neuromuscular Disord, 31, 931–942.

6. Kim, D.-S., Hayashi, Y.K., Matsumoto, H., Ogawa, M., Noguchi, S., Murakami, N., Sakuta, R., Mochizuki, M., Michele, D.E., Campbell, K.P., et al. (2004) POMT1 mutation results in defective glycosylation and loss of laminin-binding activity in &agr;-DG. Neurology, 62, 1009–1011.

7. Graziano, A., Bianco, F., D’Amico, A., Moroni, I., Messina, S., Bruno, C., Pegoraro, E., Mora, M., Astrea, G., Magri, F., et al. (2015) Prevalence of congenital muscular dystrophy in Italy: a population study. Neurology, 84, 904–911.

8. Sframeli, M., Sarkozy, A., Bertoli, M., Astrea, G., Hudson, J., Scoto, M., Mein, R., Yau, M., Phadke, R., Feng, L., et al. (2017) Congenital muscular dystrophies in the UK population: Clinical and molecular spectrum of a large cohort diagnosed over a 12-year period. Neuromuscular Disord, 27, 793–803.

9. Song, D., Dai, Y., Chen, X., Fu, X., Chang, X., Wang, N., Zhang, C., Yan, C., Zheng, H., Wu, L., et al. (2020) Genetic Variations and Clinical Spectrum of Dystroglycanopathy in a Large Cohort of Chinese Patients. Clin Genet.

10. Willer, T., Prados, B., Falcón-Pérez, J.M., Renner-Müller, I., Przemeck, G.K.H., Lommel, M., Coloma, A., Valero, M.C., Angelis, M.H. de, Tanner, W., et al. (2004) Targeted disruption of the Walker-Warburg syndrome gene Pomt1 in mouse results in embryonic lethality. Proceedings of the National Academy of Sciences of the United States of America, 101, 14126–14131.

11. Cohn, R.D., Henry, M.D., Michele, D.E., Barresi, R., Saito, F., Moore, S.A., Flanagan, J.D., Skwarchuk, M.W., Robbins, M.E., Mendell, J.R., et al. (2002) Disruption of DAG1 in differentiated skeletal muscle reveals a role for dystroglycan in muscle regeneration. Cell, 110, 639–648.

12. Satz, J.S., Barresi, R., Durbeej, M., Willer, T., Turner, A., Moore, S.A. and Campbell, K.P. (2008) Brain and eye malformations resembling Walker-Warburg syndrome are recapitulated in mice by dystroglycan deletion in the epiblast. Journal of Neuroscience, 28, 10567–10575.

13. Rubio-Fernández, M., Uribe, M.L., Vicente-Tejedor, J., Germain, F., Susín-Lara, C., Quereda, C., Montoliu, L., Villa, P. de la, Martín-Nieto, J. and Cruces, J. (2018) Impairment of photoreceptor ribbon synapses in a novel Pomt1 conditional knockout mouse model of dystroglycanopathy. Sci Rep-uk, 8, 8543.

14. Li, M., Hromowyk, K.J., Amacher, S.L. and Currie, P.D. (2017) Chapter 14 Muscular dystrophy modeling in zebrafish. Methods Cell Biol, 138, 347–380.

15. Widrick, J.J., Kawahara, G., Alexander, M.S., Beggs, A.H. and Kunkel, L.M. (2019) Discovery of Novel Therapeutics for Muscular Dystrophies using Zebrafish Phenotypic Screens. Journal of neuromuscular diseases, 6, 271–287.

16. Carss, K.J., Stevens, E., Foley, A.R., Cirak, S., Riemersma, M., Torelli, S., Hoischen, A., Willer, T., Scherpenzeel, M. van, Moore, S.A., et al. (2013) Mutations in GDP-Mannose Pyrophosphorylase B Cause Congenital and Limb-Girdle Muscular Dystrophies Associated with Hypoglycosylation of α-Dystroglycan. American journal of human genetics.

17. Stevens, E., Carss, K.J., Cirak, S., Foley, A.R., Torelli, S., Willer, T., Tambunan, D.E., Yau, S., Brodd, L., Sewry, C.A., et al. (2013) Mutations in B3GALNT2 Cause Congenital Muscular Dystrophy and Hypoglycosylation of α-Dystroglycan. American journal of human genetics.

18. Manzini, M.C., Tambunan, D.E., Hill, R.S., Yu, T.W., Maynard, T.M., Heinzen, E.L., Shianna, K.V., Stevens, C.R., Partlow, J.N., Barry, B.J., et al. (2012) Exome sequencing and functional validation in zebrafish identify GTDC2 mutations as a cause of Walker-Warburg syndrome. American journal of human genetics, 91, 541–547.

19. Costanzo, S.D., Balasubramanian, A., Pond, H.L., Rozkalne, A., Pantaleoni, C., Saredi, S., Gupta, V.A., Sunu, C.M., Yu, T.W., Kang, P.B., et al. (2014) POMK mutations disrupt muscle development leading to a spectrum of neuromuscular presentations. Human Molecular Genetics.

20. Avsar-Ban, E., Ishikawa, H., Manya, H., Watanabe, M., Akiyama, S., Miyake, H., Endo, T. and Tamaru, Y. (2010) Protein O-mannosylation is necessary for normal embryonic development in zebrafish. Glycobiology, 20, 1089–1102.

21. Gupta, V., Kawahara, G., Gundry, S.R., Chen, A.T., Lencer, W.I., Zhou, Y., Zon, L.I., Kunkel, L.M. and Beggs, A.H. (2011) The zebrafish dag1 mutant: a novel genetic model for dystroglycanopathies. Human Molecular Genetics, 20, 1712–1725.

22. Serafini, P.R., Feyder, M.J., Hightower, R.M., Garcia-Perez, D., Vieira, N.M., Lek, A., Gibbs, D.E., Moukha-Chafiq, O., Augelli-Szafran, C.E., Kawahara, G., et al. (2018) A limb-girdle muscular dystrophy 2I model of muscular dystrophy identifies corrective drug compounds for dystroglycanopathies. JCI insight, 3, 199.

23. Liu, Y., Yu, M., Shang, X., Nguyen, M.H.H., Balakrishnan, S., Sager, R. and Hu, H. (2020) Eyes shut homolog (EYS) interacts with matriglycan of O-mannosyl glycans whose deficiency results in EYS mislocalization and degeneration of photoreceptors. Sci Rep-uk, 10, 7795.

24. Liu, Y., Rittershaus, J.M., Yu, M., Sager, R. and Hu, H. (2022) Deletion of POMT2 in Zebrafish Causes Degeneration of Photoreceptors. Int J Mol Sci, 23, 14809.

25. Diesen, C., Saarinen, A., Pihko, H., Rosenlew, C., Cormand, B., Dobyns, W.B., Dieguez, J., Valanne, L., Joensuu, T. and Lehesjoki, A.-E. (2004) POMGnT1 mutation and phenotypic spectrum in muscle-eye-brain disease. Journal of Medical Genetics, 41, e115.

26. Xu, M., Yamada, T., Sun, Z., Eblimit, A., Lopez, I., Wang, F., Manya, H., Xu, S., Zhao, L., Li, Y., et al. (2016) Mutations in POMGNT1 cause non-syndromic retinitis pigmentosa. Hum Mol Genet, 25, 1479– 1488.

27. Abrams, E.W. and Mullins, M.C. (2009) Early zebrafish development: It’s in the maternal genes. Curr Opin Genet Dev, 19, 396–403.

28. Kettleborough, R.N.W., Busch-Nentwich, E.M., Harvey, S.A., Dooley, C.M., Bruijn, E. de, Eeden, F. van, Sealy, I., White, R.J., Herd, C., Nijman, I.J., et al. (2013) A systematic genome-wide analysis of zebrafish protein-coding gene function. Nature, 496, 494–497.

29. Manzini, M.C., Gleason, D., Chang, B.S., Hill, R.S., Barry, B.J., Partlow, J.N., Poduri, A., Currier, S., Galvin-Parton, P., Shapiro, L.R., et al. (2008) Ethnically diverse causes of Walker-Warburg syndrome (WWS): FCMD mutations are a more common cause of WWS outside of the Middle East. Human Mutation, 29, E231–41.

30. Godfrey, C., Clement, E., Mein, R., Brockington, M., Smith, J., Talim, B., Straub, V., Robb, S., Quinlivan, R., Feng, L., et al. (2007) Refining genotype phenotype correlations in muscular dystrophies with defective glycosylation of dystroglycan. Brain : a journal of neurology, 130, 2725–2735.

31. Renninger, S.L., Gesemann, M. and Neuhauss, S.C.F. (2011) Cone arrestin confers cone vision of high temporal resolution in zebrafish larvae. Eur J Neurosci, 33, 658–667.

32. Kohl, S., Baumann, B., Rosenberg, T., Kellner, U., Lorenz, B., Vadalà, M., Jacobson, S.G. and Wissinger, B. (2002) Mutations in the Cone Photoreceptor G-Protein α-Subunit Gene GNAT2 in Patients with Achromatopsia. Am J Hum Genetics, 71, 422–425.

33. Gao, P., Qin, Y., Qu, Z., Huang, Y., Liu, X., Li, J., Liu, F. and Liu, M. (2022) The Zpr-3 antibody recognizes the 320-354 region of Rho and labels both rods and green cones in zebrafish. Biorxiv, 2022.02.21.481375.

34. El-Brolosy, M.A. and Stainier, D.Y.R. (2017) Genetic compensation: A phenomenon in search of mechanisms. Plos Genet, 13, e1006780.

35. Rouf, M.A., Wen, L., Mahendra, Y., Wang, J., Zhang, K., Liang, S., Wang, Y., Li, Z., Wang, Y. and Wang, G. (2022) The Recent Advances and Future Perspectives of Genetic Compensation Studies in the Zebrafish Model. Genes Dis.

36. White, R.J., Collins, J.E., Sealy, I.M., Wali, N., Dooley, C.M., Digby, Z., Stemple, D.L., Murphy, D.N., Billis, K., Hourlier, T., et al. (2017) A high-resolution mRNA expression time course of embryonic development in zebrafish. Elife, 6, e30860.

37. Harvey, S.A., Sealy, I., Kettleborough, R., Fenyes, F., White, R., Stemple, D. and Smith, J.C. (2013) Identification of the zebrafish maternal and paternal transcriptomes. Development, 140, 2703–2710.

38. Novak, J.S., Spathis, R., Dang, U.J., Fiorillo, A.A., Hindupur, R., Tully, C.B., Mázala, D.A.G., Canessa, E., Brown, K.J., Partridge, T.A., et al. (2021) Interrogation of Dystrophin and Dystroglycan Complex Protein Turnover After Exon Skipping Therapy. J Neuromuscul Dis, 8, S383–S402.

39. Andersson, M. and Kettunen, P. (2021) Effects of Holding Density on the Welfare of Zebrafish: A Systematic Review. Zebrafish, 18, 297–306.

40. El-Brolosy, M.A., Kontarakis, Z., Rossi, A., Kuenne, C., Günther, S., Fukuda, N., Kikhi, K., Boezio, G.L.M., Takacs, C.M., Lai, S.-L., et al. (2019) Genetic compensation triggered by mutant mRNA degradation. Nature, 568, 193–197.

41. Bai, L., Kovach, A., You, Q., Kenny, A. and Li, H. (2019) Structure of the eukaryotic protein O-mannosyltransferase Pmt1–Pmt2 complex. Nat Struct Mol Biol, 26, 704–711.

42. Kawahara, G., Guyon, J.R., Nakamura, Y. and Kunkel, L.M. (2010) Zebrafish models for human FKRP muscular dystrophies. Human Molecular Genetics, 19, 623–633.

43. Gupta, V.A., Kawahara, G., Myers, J.A., Chen, A.T., Hall, T.E., Manzini, M.C., Currie, P.D., Zhou, Y., Zon, L.I., Kunkel, L.M., et al. (2012) A splice site mutation in laminin-α2 results in a severe muscular dystrophy and growth abnormalities in zebrafish. PLoS ONE, 7, e43794.

44. Tonelotto, V., Consorti, C., Facchinello, N., Trapani, V., Sabatelli, P., Giraudo, C., Spizzotin, M., Cescon, M., Bertolucci, C. and Bonaldo, P. (2022) Collagen VI ablation in zebrafish causes neuromuscular defects during developmental and adult stages. Matrix Biol, 112, 39–61.

45. Clements, R., Turk, R., Campbell, K.P. and Wright, K.M. (2017) Dystroglycan Maintains Inner Limiting Membrane Integrity to Coordinate Retinal Development. Journal of Neuroscience, 37, 8559–8574.

46. Lindenmaier, L.B., Parmentier, N., Guo, C., Tissir, F. and Wright, K.M. (2019) Dystroglycan is a scaffold for extracellular axon guidance decisions. eLife, 8, 239.

47. Clements, R. and Wright, K.M. (2018) Retinal ganglion cell axon sorting at the optic chiasm requires dystroglycan. Dev Biol, 442, 210–219.

48. Wright, K.M., Lyon, K.A., Leung, H., Leahy, D.J., Ma, L. and Ginty, D.D. (2012) Dystroglycan organizes axon guidance cue localization and axonal pathfinding. Neuron, 76, 931–944.

49. Davison, C. and Zolessi, F.R. (2021) Slit2 is necessary for optic axon organization in the zebrafish ventral midline. Cells Dev, 166, 203677.

50. Laue, K., Rajshekar, S., Courtney, A.J., Lewis, Z.A. and Goll, M.G. (2019) The maternal to zygotic transition regulates genome-wide heterochromatin establishment in the zebrafish embryo. Nat. Commun., 10, 1551.

51. Han, R., Kanagawa, M., Yoshida-Moriguchi, T., Rader, E.P., Ng, R.A., Michele, D.E., Muirhead, D.E., Kunz, S., Moore, S.A., Iannaccone, S.T., et al. (2009) Basal lamina strengthens cell membrane integrity via the laminin G domain-binding motif of α-dystroglycan. Proc. Natl. Acad. Sci., 106, 12573–12579.

52. Franekova, V., Storjord, H.I., Leivseth, G. and Nilssen, Ø. (2021) Protein homeostasis in LGMDR9 (LGMD2I) – The role of ubiquitin–proteasome and autophagy–lysosomal system. Neuropathol. Appl. Neurobiol., 47, 519–531.

53. Bailey, E.C., Alrowaished, S.S., Kilroy, E.A., Crooks, E.S., Drinkert, D.M., Karunasiri, C.M., Belanger, J.J., Khalil, A., Kelley, J.B. and Henry, C.A. (2019) NAD+ improves neuromuscular development in a zebrafish model of FKRP-associated dystroglycanopathy. Skeletal muscle, 9, 21–23.

54. Gee, S.H., Montanaro, F., Lindenbaum, M.H. and Carbonetto, S. (1994) Dystroglycan-α, a dystrophin-associated glycoprotein, is a functional agrin receptor. Cell, 77, 675–686.

55. Jacobson, C., Côté, P.D., Rossi, S.G., Rotundo, R.L. and Carbonetto, S. (2001) The Dystroglycan Complex Is Necessary for Stabilization of Acetylcholine Receptor Clusters at Neuromuscular Junctions and Formation of the Synaptic Basement Membrane. J Cell Biology, 152, 435–450.

56. Sheikh, M.O., Capicciotti, C.J., Liu, L., Praissman, J., Ding, D., Mead, D.G., Brindley, M.A., Willer, T., Campbell, K.P., Moremen, K.W., et al. (2022) Cell surface glycan engineering reveals that matriglycan alone can recapitulate dystroglycan binding and function. Nat Commun, 13, 3617.

57. Sato, S., Omori, Y., Katoh, K., Kondo, M., Kanagawa, M., Miyata, K., Funabiki, K., Koyasu, T., Kajimura, N., Miyoshi, T., et al. (2008) Pikachurin, a dystroglycan ligand, is essential for photoreceptor ribbon synapse formation. Nature Neuroscience, 11, 923–931.

58. Plump, A.S., Erskine, L., Sabatier, C., Brose, K., Epstein, C.J., Goodman, C.S., Mason, C.A. and Tessier-Lavigne, M. (2002) Slit1 and Slit2 cooperate to prevent premature midline crossing of retinal axons in the mouse visual system. Neuron, 33, 219–232.

59. Davison, C. and Zolessi, F.R. (2021) Slit2 is necessary for optic axon organization in the zebrafish ventral midline. Cells Dev, 166, 203677.

60. Omori, Y., Araki, F., Chaya, T., Kajimura, N., Irie, S., Terada, K., Muranishi, Y., Tsujii, T., Ueno, S., Koyasu, T., et al. (2012) Presynaptic Dystroglycan-Pikachurin Complex Regulates the Proper Synaptic Connection between Retinal Photoreceptor and Bipolar Cells. J Neurosci, 32, 6126–6137.

61. Garcia-Delgado, A.B., Valdes-Sanchez, L., Morillo-Sanchez, M.J., Ponte-Zuñiga, B., Diaz-Corrales, F.J. and Cerda, B. de la (2021) Dissecting the role of EYS in retinal degeneration: clinical and molecular aspects and its implications for future therapy. Orphanet J. Rare Dis., 16, 222.

62. Kawahara, G., Gasperini, M.J., Myers, J.A., Widrick, J.J., Eran, A., Serafini, P.R., Alexander, M.S., Pletcher, M.T., Morris, C.A. and Kunkel, L.M. (2014) Dystrophic muscle improvement in zebrafish via increased heme oxygenase signaling. Human Molecular Genetics, 23, 1869–1878.

63. Schindelin, J., Arg, I., Arganda-Carreras, I., a-Carreras, Frise, E., Kaynig, V., Longair, M., Pietzsch, T., Preibisch, S., Rueden, C., et al. (2012) Fiji: an open-source platform for biological-image analysis. 9, 676–82.

64. Stirling, D.R., Swain-Bowden, M.J., Lucas, A.M., Carpenter, A.E., Cimini, B.A. and Goodman, A. (2021)yyy CellProfiler 4: improvements in speed, utility and usability. Bmc Bioinformatics, 22, 433.

