## Supplemental Figures for "Removal of *pomt1* in zebrafish leads to loss of α-dystroglycan glycosylation and dystroglycanopathy phenotypes"

Brittany F. Karas, et al.

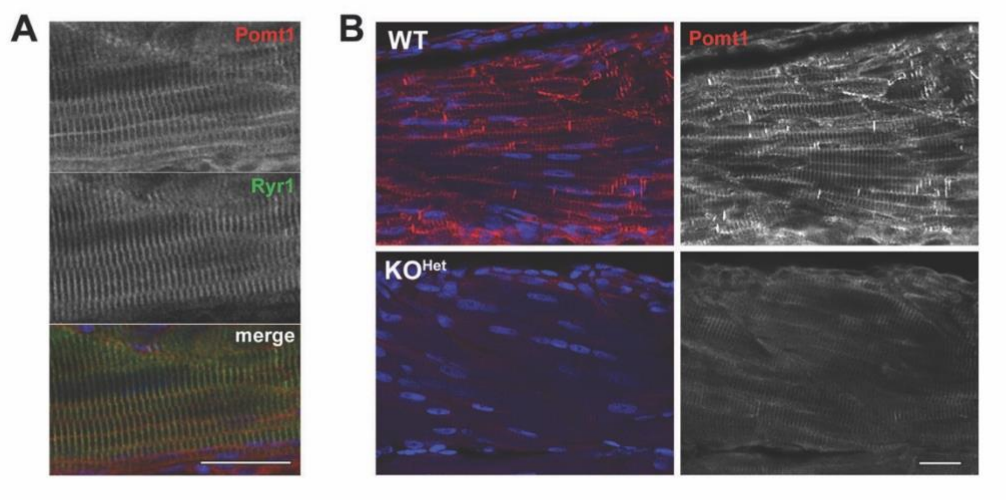


**Supplementary Figure 1. Pomt1 antibody validation and KO analysis. A.** Immunostaining for ryanodine receptor (Ryr1) was used to outline the sarcoplasmic reticulum and show Pomt1 localization. Scale bar: 20 $\mu$m. **B.** Immunohistochemistry of cryosections from 10 dpf muscle shows loss of Pomt1 in $KO^{Het}$. Scale bar: 20 $\mu$m.

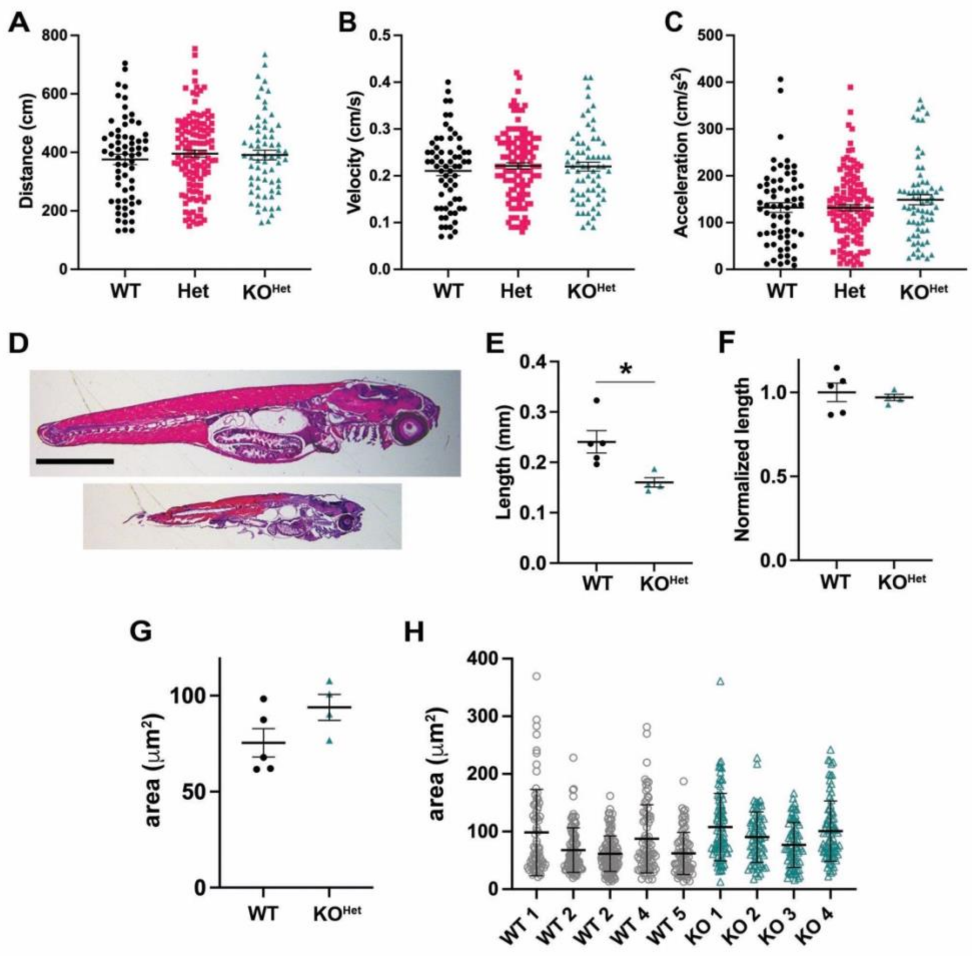


**Supplementary Figure 2. Locomotor analysis in *pomt1* KO$^{Het}$ larvae at 5 dpf. A-C.** No differences from WT and Het were found in mobility assays in 5 dpf *pomt1* KO$^{Het}$ larvae in distance traveled (**A.**), velocity (**B.**) and acceleration (**C.**). Larvae tested: WT = 63, Het = 122, KO$^{Het}$ = 57, N=3 clutches. **D-F.** Myofiber length at 30 dpf measured as the distance across myosepta/MTJs in histological sections stained with H&E (**D.** Scale bar: 2mm) was shorter (**E.** Length: WT 240.6±22.1 μm n=5 fish, KO$^{Het}$ 160.3±9.3 μm n=5 fish, p=0.019 *) but it was not changed when the length was normalized to the length of the fish (**F.** Normalized length: WT 1.00±0.05 n=5 fish, KO$^{Het}$ 0.97±0.01 n=5 fish, p=0.658) **F. G.** Fiber area in dorsal muscle from 30 dpf dorsal muscles from 5 independent juveniles was not significantly changed (Area: WT 75.4±7.4 μm$^2$ n=5 fish, KO$^{Het}$ 90.6±6.2 μm$^2$ n=5 fish, p=0.156). **H.** Fiber size distribution for the experimental animals.

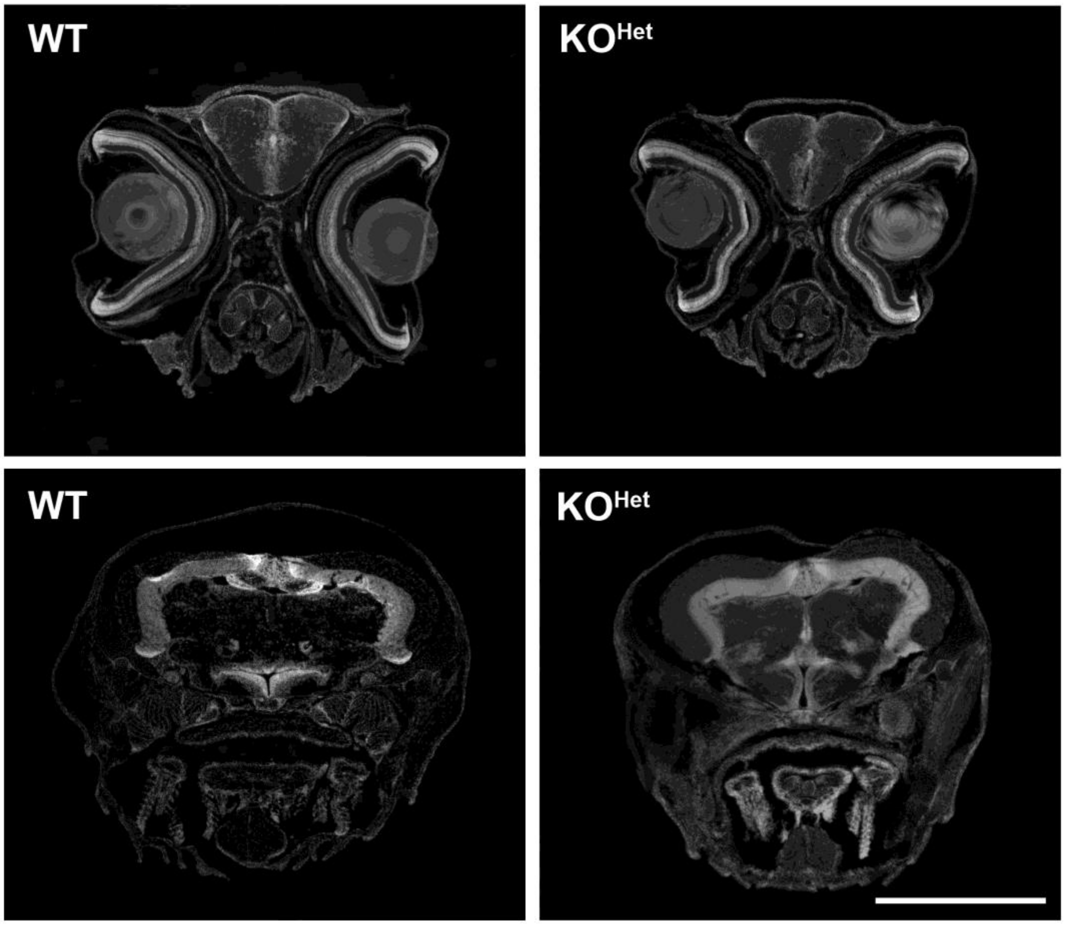


**Supplementary Figure 3. Brain histology of *pomt1* KO$^{Het}$ juveniles at 30 dpf.** No differences in brain structure or hydrocephaly are observed at 30 dpf in *pomt1* KO$^{Het}$ juveniles. Representative images from DAPI stained axial cryosections are shown at the level of the telencephalon (top row) and of the mesencephalon and optic tectum (bottom row). Scale bar: 400 μm

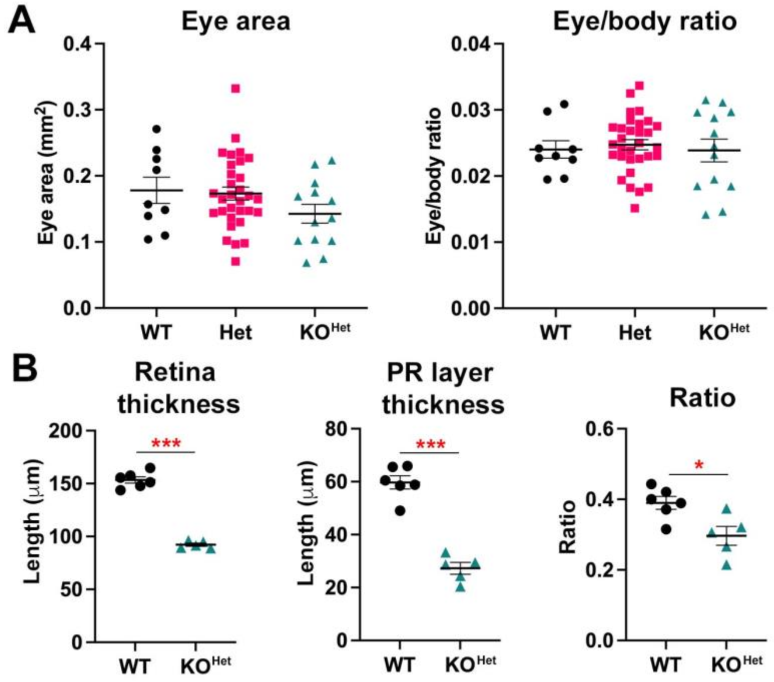


**Supplementary Figure 4. Eye and retinal measurements in 30 dpf *pomt1* fish. A.** Eye area (left graph) showed a trend towards being reduced but was highly variable with no significant differences (WT: 0.178±0.020 mm$^2$, Het: 0.173±0.001, KO: 0.142±0.014). When eye size was normalized to body size (right graph), there was no difference in the eye/body ratio showing that the eye size was proportional to the body (WT: 0.024±0.0013, Het: 0.025±0.0001, KO: 0.024±0.0017; WT n=9, Het n=31, KO n=13, N=2 clutches). **B.** Retinal thickness (left) and photoreceptor (PR) layer thickness measures on a subset of small WT (n=6) and KO (n=5) animals showed a significant reduction (Retinal thickness. WT: 153.5±3.1μm, KO: 92.3±1.5μm, p<0.0001 ***. PR layer thickness. WT: 59.7±2.5μm, KO: 27.3±2.2μm, p<0.0001 ***). The PR layer was disproportionally reduced when compared to total retinal thickness (WT: 0.390±0.018, KO: 0.297±0.027, p=0.015 *).

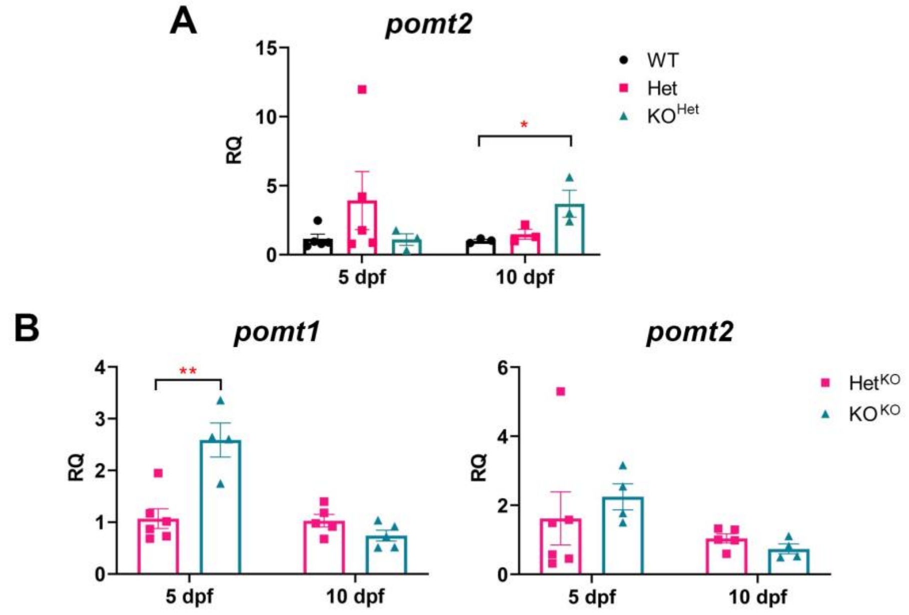


**Supplementary Figure 5. Expression analysis of *pomt2* in Het X Het and KO X Het crosses. A.** *pomt2* showed a trend for increased expression by qPCR in Het larvae at 5 dpf and an increase in KO$^{Het}$ larvae at 10 dpf. This latest timepoint matches the increase in *pomt1* expression noted in **Fig.1C. B.** *pomt1* expression showed an increase at 5dpf in KO$^{KO}$ larvae, but then declined by 10 dpf (left panel). *pomt2* showed no differences at either timepoint (right panel).

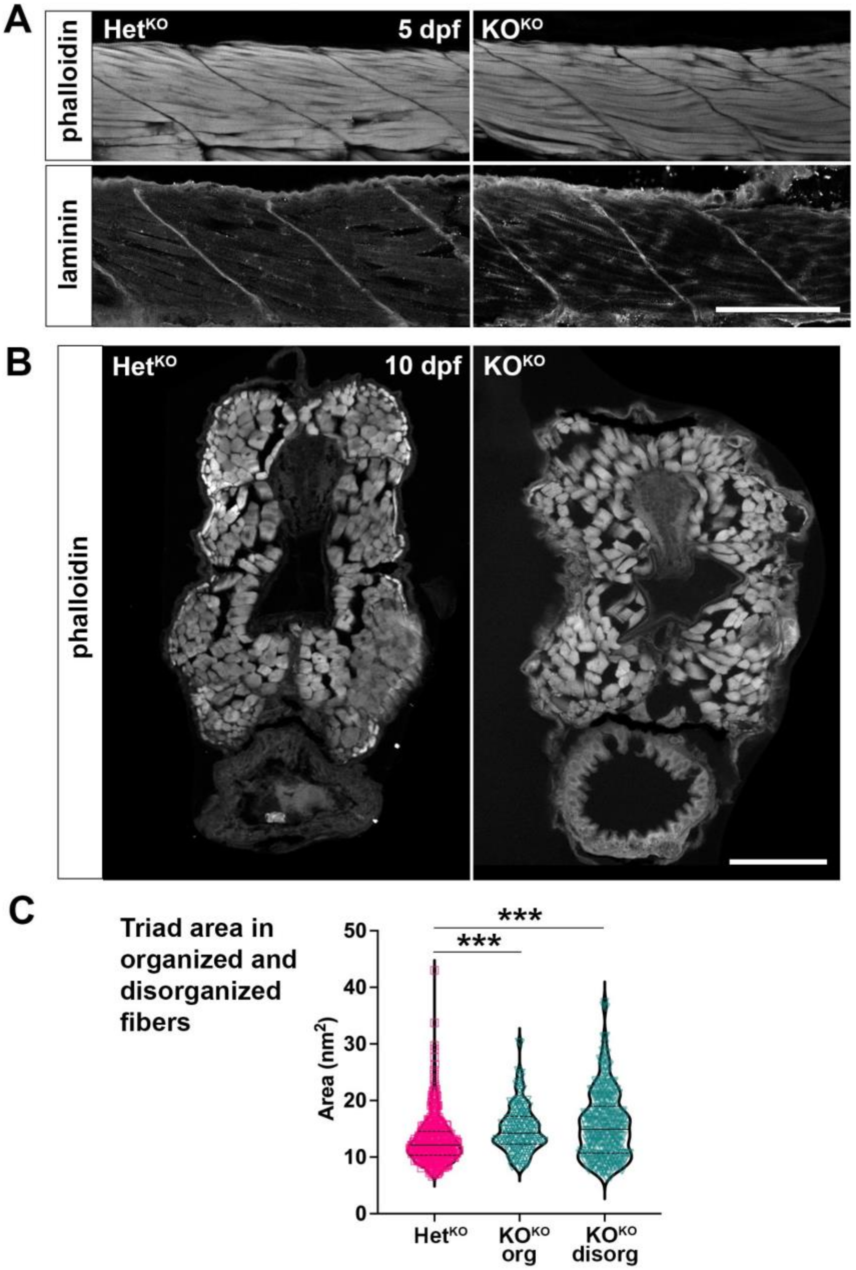


**Supplementary Figure 6. Muscle integrity analysis in *pomt1* KO$^{KO}$ larvae at 5 dpf. A.** Muscle integrity looks comparable at 5 dpf in Het$^{KO}$ and KO$^{KO}$ larvae upon staining with phalloidin to show actin filaments and $\alpha$-bungarotoxin to identify the neuromuscular junction. Scale bar: 100 $\mu$m. **B.** Muscle fiber disorganization is also observed in 10 dpf KO$^{KO}$ muscle in axial sections stained for phalloidin. Scale bar: 50 $\mu$m. **C.** Quantification of triad area sorted by organized/aligned (org) and disorganized/misaligned (disorg) muscle fibers as shown in **Fig.7A**. Triad size is increased in both, but much more variable in disorganized fibers (Fiber area: Het$^{KO}$ 12.9±0.2 $\mu m^2$ n=387 triads, org KO$^{KO}$ 14.9±0.3 $\mu m^2$ n=174 triads, disorg KO$^{KO}$ 15.7±0.4 $\mu m^2$ n=261 triads, p<0.0001 ***).

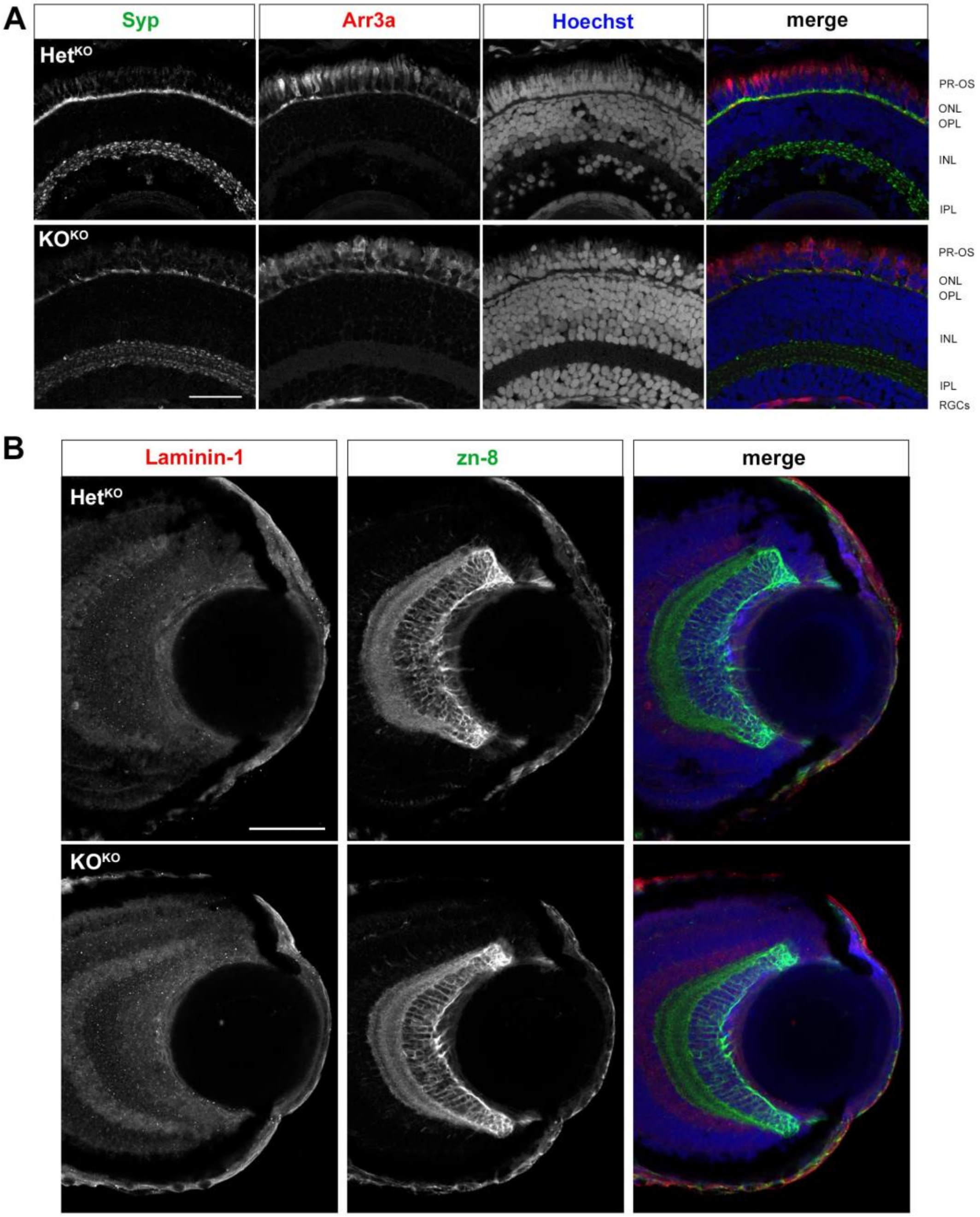


**Supplementary Figure 7. Additional imaging of the 5 dpf *pomt1* KO$^{KO}$ retina. A.** Larger field from 40X magnification images used in **Figure 8A** showing discontinuities in synaptophysin (Syp) staining and nuclear disorganization in the photoreceptor layer stained with Hoechst in the KO$^{KO}$ retina. Scale bar: 30 μm. **B.** Laminin-1 staining outlines the basement membrane around the eye, but only shows limited staining in the inner limiting membrane at the interface of the lens and the retinal ganglion cell layer. Retinal ganglion cells are labels using zn8. Scale bar: 50 μm

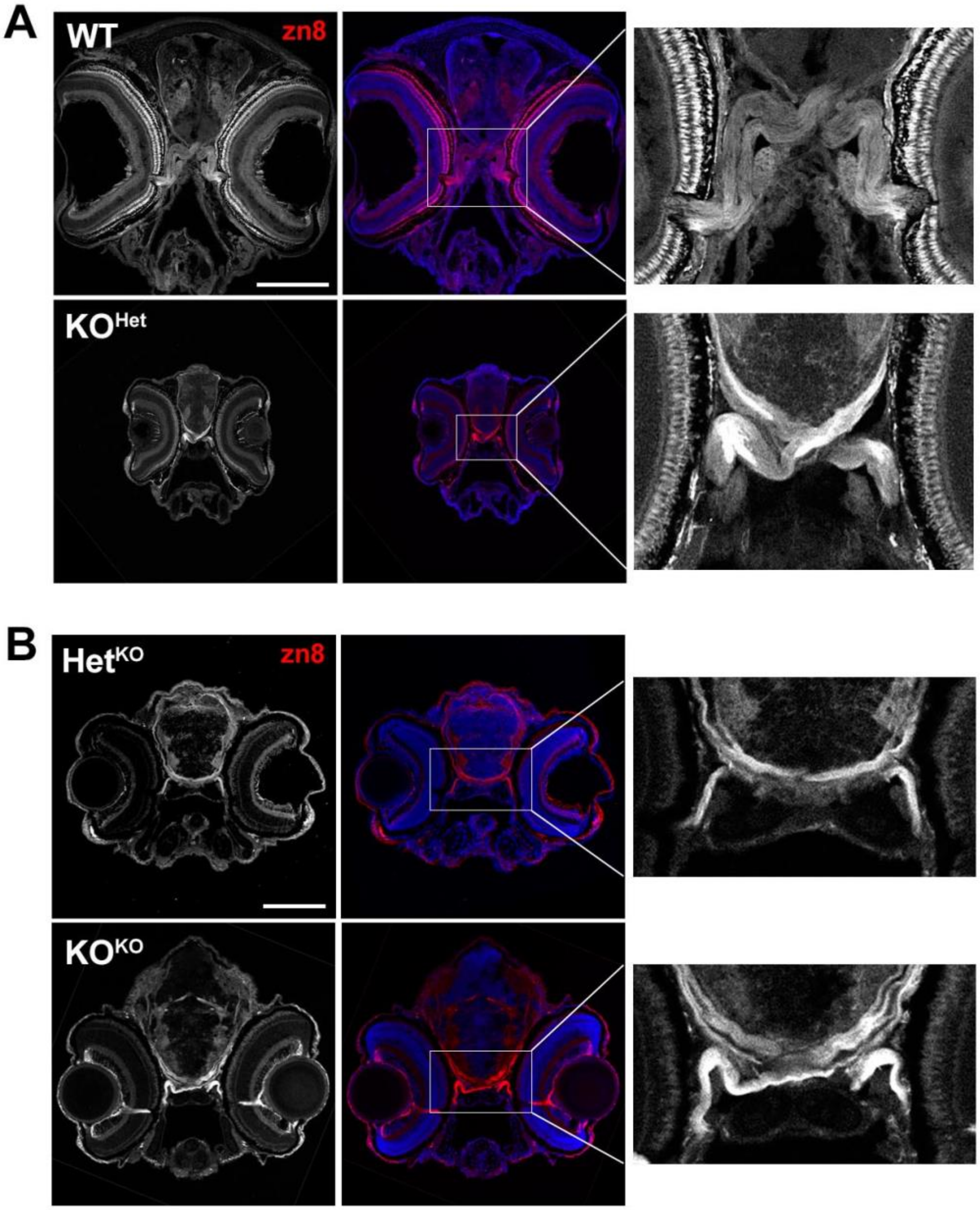


**Supplementary Figure 8. Optic chiasm axon crossing and morphology in *pomt1* KO$^{Het}$ 30 dpf juveniles and KO$^{KO}$ 5 dpf larvae. A.** While overall smaller 30 dpf *pomt1* KO$^{Het}$ brain sections stained with the neuronal marker zn8 that labels retinal ganglion cells and their axons show comparable optic chiasm organization and optic nerve structure. Scale bar: 300 μm. **B.** Corresponding whole head images showing the location of the optic chiasm shown in **Figure 8C**. Scale bar: 100 μm
